# Molecular basis of ligand binding and receptor activation at the human A3 adenosine receptor

**DOI:** 10.1101/2025.01.12.632629

**Authors:** Liudi Zhang, Jesse I. Mobbs, Felix M. Bennetts, Hariprasad Venugopal, Anh T.N. Nguyen, Arthur Christopoulos, Daan van der Es, Laura H. Heitman, Lauren T. May, Alisa Glukhova, David M. Thal

## Abstract

Adenosine receptors (ARs: A_1_AR, A_2A_AR, A_2B_AR, and A_3_AR) are crucial therapeutic targets, yet developing selective, efficacious drugs remains challenging. Here, we present high-resolution cryo-electron microscopy (cryo-EM) structures of the human A_3_AR in three distinct functional states: bound to the endogenous agonist adenosine, the clinically relevant agonist Piclidenoson, and the covalent antagonist LUF7602. These structures, complemented by mutagenesis and pharmacological studies, reveal a unique A_3_AR activation mechanism involving an extensive hydrogen bond network from the extracellular surface down to the orthosteric binding site. In addition, we identify a cryptic pocket that accommodates the N^6^-iodobenzyl group of Piclidenoson through a ligand-dependent conformational change of M174^5.35^. Our comprehensive structural and functional characterization of A_3_AR advances understanding of adenosine receptor pharmacology and establishes a foundation for developing more selective therapeutics for various disorders including inflammatory diseases, cancer, and glaucoma.

**Teaser:** Structures of the A_3_AR in different conformations reveal a unique activation mechanism and cryptic binding pocket.

## Introduction

The four adenosine receptors (ARs: A_1_AR, A_2A_AR, A_2B_AR, and A_3_AR) are Class A G protein-coupled receptors (GPCRs) activated by extracellular levels of the nucleoside adenosine (*1*). These receptors are broadly expressed in humans, and they regulate a diverse range of physiological processes. Tremendous effort has gone into modulating the activity of adenosine receptors as potential treatments for cardiovascular disease, nervous system disorders, inflammation, renal and endocrine disorders, cancer, and visual disorders (*1–3*). In this regard, the human A_3_AR is enigmatic because it plays dual roles under different pathophysiological conditions (*4, 5*). This complexity is particularly evident in cancer biology. For example, A_3_AR is overexpressed in several types of tumours and is a proposed diagnostic marker (*6–9*). A_3_AR overexpression suggests a pro-tumoral role, promoting cell proliferation and survival (*10–12*), while in other cancer types, the activation of A_3_AR demonstrated anti-tumoral effects by triggering apoptosis and inhibiting cell growth (*13–15*). Nevertheless, the A_3_AR selective agonists Piclidenoson and Namodenoson have progressed into clinical trials for treating inflammatory diseases, including rheumatoid arthritis, psoriasis, and liver diseases such as hepatocellular carcinoma, hepatitis, and dry eye syndrome (*16–22*). In contrast, A_3_AR antagonists are being developed as treatments for glaucoma and asthma (*23–26*).

Despite the vast therapeutic potential that adenosine receptors hold, few drug candidates have progressed through the clinic. A major challenge in GPCR drug discovery, particularly relevant to adenosine receptors, is identifying ligands that selectively target one receptor subtype over similarly related subtypes; lack of such selectivity can lead to off-target side effects. To address this challenge, structural studies of GPCRs with different types of ligands have begun to illuminate molecular mechanisms of ligand selectivity, or lack thereof, paving the way for a new era of drug discovery. Regarding adenosine receptors, the A_2A_AR was a model GPCR for pioneering structural biology work (*27*) with various antagonist, agonist, and partial agonist-bound structures available, including the receptor in inactive (*28*), intermediate (*29, 30*), and fully active conformations (*31*). Structures of the A_1A_R, A_2A_AR, and A_2B_AR have revealed key principles of adenosine receptor activation and selectivity that have guided drug design efforts (*32–36*). Recent experimental structures of the A_3_AR have been determined with the ligands Piclidenoson and Namodenoson (*37, 38*). However, crucially, these structures are relatively low resolution and important parts of the ligand, an N^6^-iodobenzyl group, were not adequately resolved in the structures. Moreover, structures of the A_3_AR in the inactive and active conformations are crucial for elucidating its activation mechanism and for guiding the design of future subtype-selective ligands.

The search for potent and selective A_3_AR antagonists led to the discovery of LUF7602 (*39*), which is a covalent antagonist that was designed based on a high-affinity tricyclic xanthine scaffold (*40*) and the A_1_AR covalent antagonist FSCPX (*41*). We determined a Fab-assisted (*42, 43*) cryo-EM structure of the human A_3_AR in the inactive conformation, to a resolution of 2.8 Å, in complex with the covalent antagonist LUF7602. In addition, we report cryo-EM structures of the human wild-type A_3_AR in complex with G protein and bound to the endogenous agonist adenosine and the clinically relevant agonist Piclidenoson.

Our A_3_AR structures, combined with pharmacological data, provide insights into the ligand binding modes of both antagonists and agonists. Comparison of the inactive and active conformations of the human A_3_AR revealed a molecular basis for receptor activation. This activation is mediated by an extensive hydrogen bond network that extends from the top of TM7 through TM1, TM2, and TM3 down to the core of the orthosteric binding site. These findings not only enhance our understanding of A_3_AR ligand binding, activation, and signalling mechanisms but also provide a robust structural framework for the rational design of highly selective A_3_AR ligands that could lead to new treatment strategies for a wide range of disorders, including inflammatory diseases, cancer, and glaucoma.

## Results

### Structures of the inactive conformations of the human A_3_AR

Expression of human A_3_AR was enhanced by inserting the first 22 amino acids from the human M_4_ muscarinic receptor (M_4_ mAChR) between an N-terminal FLAG epitope and the wild-type (WT) human A_3_AR sequence **(Figure S1A)** (*32*). The pharmacological behaviour of the A_3_AR construct was assessed using a [^35^S]-GTPγS binding assay and showed minimal differences in receptor activation mediated by the agonist NECA compared to WT A_3_AR **(Figure S1B)**. To determine the inactive state structure of the A_3_AR using cryo-EM, we introduced several modifications to improve protein stability and expression (**Figure S2A**). The intracellular loop 3 (ICL3) was replaced with BRIL, and the S97R^3.39^ mutation was introduced to stabilize the inactive conformation (*44–46*). Additionally, we removed a potential glycosylation site in ECL2 (N160A) to reduce conformational heterogeneity (*47*). Residue numbering follows the numbering scheme by Ballesteros and Weinstein (*48*) and the GPCRdb numbering scheme for the ECL regions (*49*).

Purified A_3_AR-BRIL-S97R was incubated with an anti-BRIL Fab fragment (BAG2) and an anti-BAG2 nanobody (Nb) to facilitate structure determination, serving as fiducial markers and increasing the molecular weight of the complex **(Figure S2B-F)** (*42, 43*). We initially attempted to determine the A_3_AR structure with the antagonist MRS1220 (*50*). The global resolution reached 3.7 Å, with well-defined regions for the Fab-Nb-BRIL complex and lower TM segments of A_3_AR **(Figure S3)**. However, no cryo-EM density was observed in the orthosteric site, suggesting the structure was either ligand-free or had low ligand occupancy **(Figure S3E,F)**. Additionally, the cryo-EM density corresponding to ECL2 was too ambiguous for accurate modelling, though ECL1 and ECL3 backbones were traceable **(Figure S3D)**. To improve resolution, we used the covalent antagonist LUF7602 (**Figure 1A**).

**Fig. 1.**
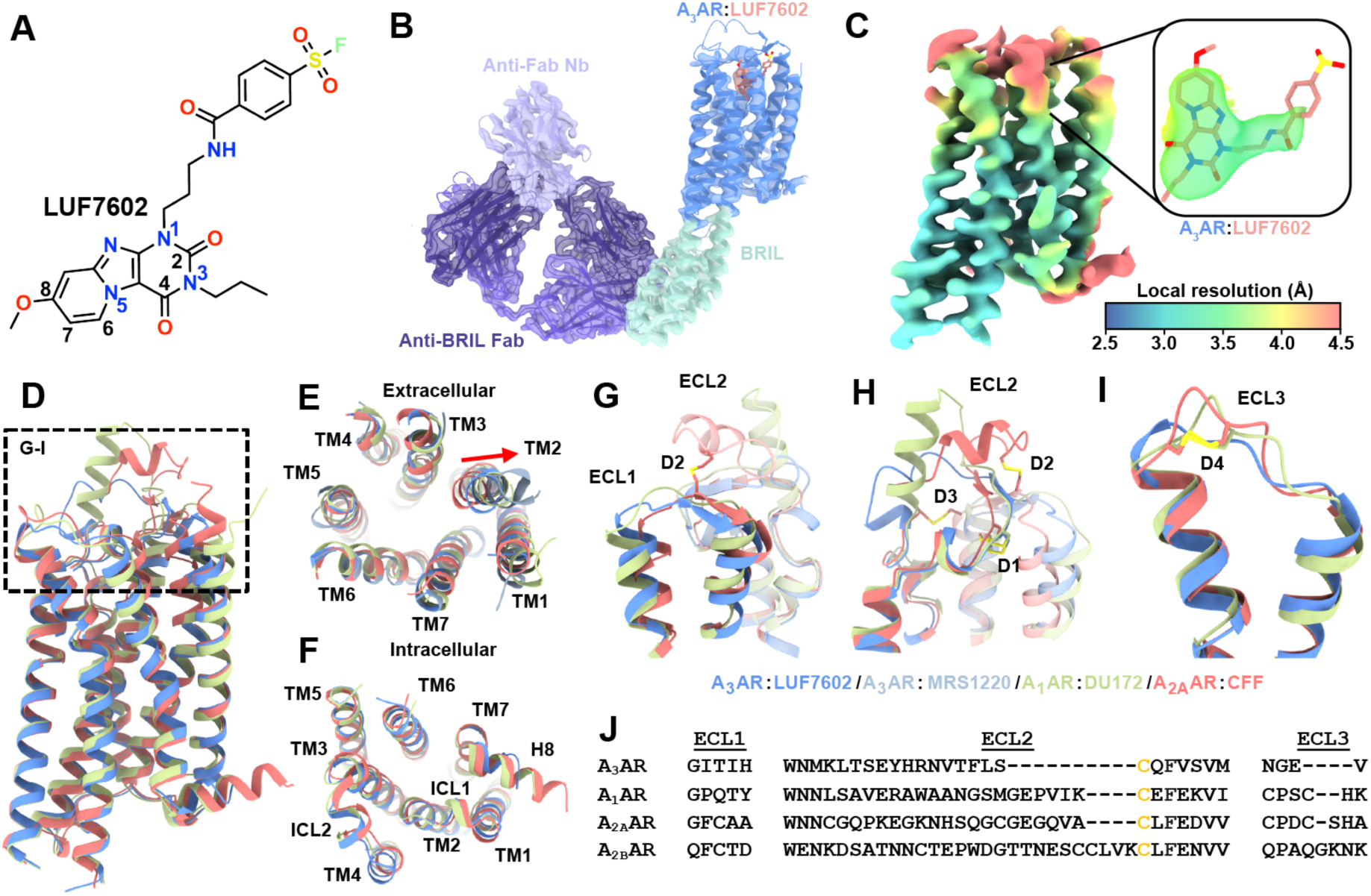
Cryo-EM structure of the A_3_AR bound to LUF7602 and comparison of the inactive conformation of adenosine receptors. **(A)** Chemical structure of LUF7602 with key atoms numbered. **(B)** Cryo-EM density map of the full inactive A_3_AR:LUF7602 complex, showing the receptor (blue), LUF7602 (peach), BRIL fusion protein (light green), anti-Fab nanobody (light purple), and anti-BRIL Fab (dark purple). **(C)** Local resolution receptor focused cryo-EM map with inset of cryo-EM density around LUF7602. **(D)** Overall structural alignment of inactive A_3_AR:LUF7602 (blue), A_1_AR:DU172 (PDB: 5UEN, light green), and A_2A_AR:CFF (PDB: 5MZP, peach). **(E)** Extracellular and **(F)** intracellular views of the aligned structures, showing transmembrane helices (TM1-TM7) and helix 8 (H8). **(G-I)** Detailed views of extracellular loops (G) ECL1, (H) ECL2, (I) ECL3 and associated helices with disulphide bonds (D1-D4) shown as sticks. **(J)** Sequence alignment of ECL1, ECL2, and ECL3 for the adenosine receptors.

A NanoBRET binding assay (*51, 52*) using an N-terminal NanoLuc (Nluc-A_3_AR) tagged receptor and the fluorescent antagonist XAC analog (XAC-630) (*53*) confirmed that LUF7602 exhibited wash-resistant inhibition with consistent binding affinity across A_3_AR-WT, A_3_AR-BRIL, and A_3_AR-BRIL-S97R constructs **(Figure S2G-H)** (*39, 54*). We determined the structure of the A_3_AR-LUF7602 complex at a global resolution of 2.7 Å (**Figure 1B, S4, Table S1)**. Focused refinement further improved the quality of the receptor density yielding a map at 3.3 Å (**Figure 1C, S4)**, enabling clear assignment of most receptor residues except for those near the covalent sulfonyl group attachment and Y265^7.36^ (**Figure 1C, Figure S5A)**.

### Comparison of the inactive conformation of adenosine receptors

The overall inactive conformation of LUF7602-bound A_3_AR shares similarities with other adenosine receptor structures bound to xanthine-based antagonists, including structures of the A_2A_AR bound to caffeine (PDB: 5MZP), theophylline (PDB: 5MZJ), XAC (PDB: 3REY), Istradefylline (PDB: 8GNG) (*33, 55, 56*), and the A_1_AR bound to the irreversible antagonist DU172 (PDB: 5UEN) and PSB36 (PDB: 5N2S) (**Figure 1D**) (*32, 33*). Despite relatively low sequence similarity, globally, the inactive conformation of the A_1_AR, A_2A_AR, and A_3_AR receptors align well with RMSD values of less than 1 Å across the TM regions (**Figure 1D-J**). Regions of high similarity include the positions of the TM helices (**Figure 1D**), the intracellular loops (ICLs) (**Figure 1F**), and the C-terminal region of ECL2, which is constrained due to a conserved disulphide bond (D1), C83^3.25^ and C166^ECL2^, that is common in Class A GPCRs (**Figure 1H**).

Divergence in the inactive conformation of adenosine receptor structures arises in the extracellular regions of the TM helices and ECLs (**Figure 1E-J**). These regions have the lowest sequence similarity across subtypes, with ECL2 and ECL3 varying in length. Indeed, the ECLs have been demonstrated to play a pivotal role in facilitating ligand entry and binding at adenosine receptors and other GPCRs (*27, 57–63*). The conformation of ECL1 was similar across adenosine receptor subtypes, with the C-terminal region of ECL1 forming interactions that stabilise the conformation of ECL2 (**Figure 1G**). There is more divergence near the TM2 region of ECL2. For example, compared to the A_2A_AR, there was a 5Å outward displacement of TM2 in the DU172-bound A_1_AR structure due to the proximity of the benzene-sulfonate linkage that covalently links DU172 to TM7. A similar 3Å TM2 shift was observed in the LUF7602-bound A_3_AR structure. Interestingly, a comparison with our MRS1220-A_3_AR dataset revealed an 8 Å outward shift of TM2. However, the overall significance of this is caveated by the low resolution of the structure (**Figure 1E**, light blue). Nevertheless, the different conformations of TM2 and ECL1 highlight their role in ligand binding.

The structure of ECL2 varies across AR subtypes, with differences in the number of disulphide bonds in each AR subtype affecting the tertiary loop structure (with 1 ECL2 disulphide bond in the A_1_AR and A_3_AR, 2 in A_2B_AR, and 3 in A_2A_AR) (**Figure 1H**). In the A_1_AR, ECL2 forms a 3-turn α-helix that extends perpendicular to the membrane and then loops down to form an anti-parallel β-strand with ECL1, extending over the orthosteric binding site towards TM5 (**Figure 1H).** In contrast, even though the length of ECL2 in the A_2A_AR is similar to the A_1_AR, the loop coils upward, forming a disulphide bond with TM3 followed by a 2.5-turn α-helix parallel to the membrane that also forms a disulphide bond with ECL1 before looping across the orthosteric site. The A_3_AR has the shortest ECL2, with a small helix parallel to the membrane, before looping over to an anti-parallel β-strand with ECL1 followed by the conserved α-helix over the orthosteric site. The ECL2 of the A_2B_AR is the longest in sequence, but structural information for this loop is lacking. The conformation of ECL3 appears to be similar across adenosine receptor subtypes, except for the A_3_AR lacking an initial loop following TM6 due to the lack of a disulphide bond and a shorter sequence (**Figure 1I**).

### Ligand interactions at the inactive A_3_AR

Clear EM density revealed LUF7602 deeply bound in the A_3_AR orthosteric site, interacting with TM helices 1-3, 5-7, and ECL2 (**Figure 2A,B**). The xanthine core of LUF7602 formed a π-π stacking interaction with F168^45.52^, a common adenosine receptor interaction (**Figure 2C**). The covalent attachment of LUF7602 occured via its benzene-sulfonate group and Y265^7.36^ (**Figure 2D**). To probe the pharmacology of the A_3_AR orthosteric binding site and validate observed interactions from the ligand-bound A_3_AR structures, we used a saturation binding NanoBRET assay to measure the binding affinity of the fluorescent antagonist XAC-630 to wild-type (WT) and mutant A_3_AR constructs (**Figure 2G,H**, **Table 1**). Mutations were based on residues interacting with LUF7602, adenosine, and Piclidenoson. XAC-630 showed no specific binding to the N250A^6.55^ and H272A^7.43^ mutants, consistent with previous findings (*64*). Overall, the point mutations had little effect on the binding affinity of XAC-630 with the H95A^3.37^ and H95F^3.37^ mutations reducing affinity ∼5-fold and the Y15A^1.35^ and M174A^5.35.^ mutations reducing affinity 3-fold. However, multiple mutations significantly decreased B_max_ values compared to WT A_3_AR indicating these residues lowered receptor expression (**Figure 2G**, **Table 1**).

**Fig. 2.**
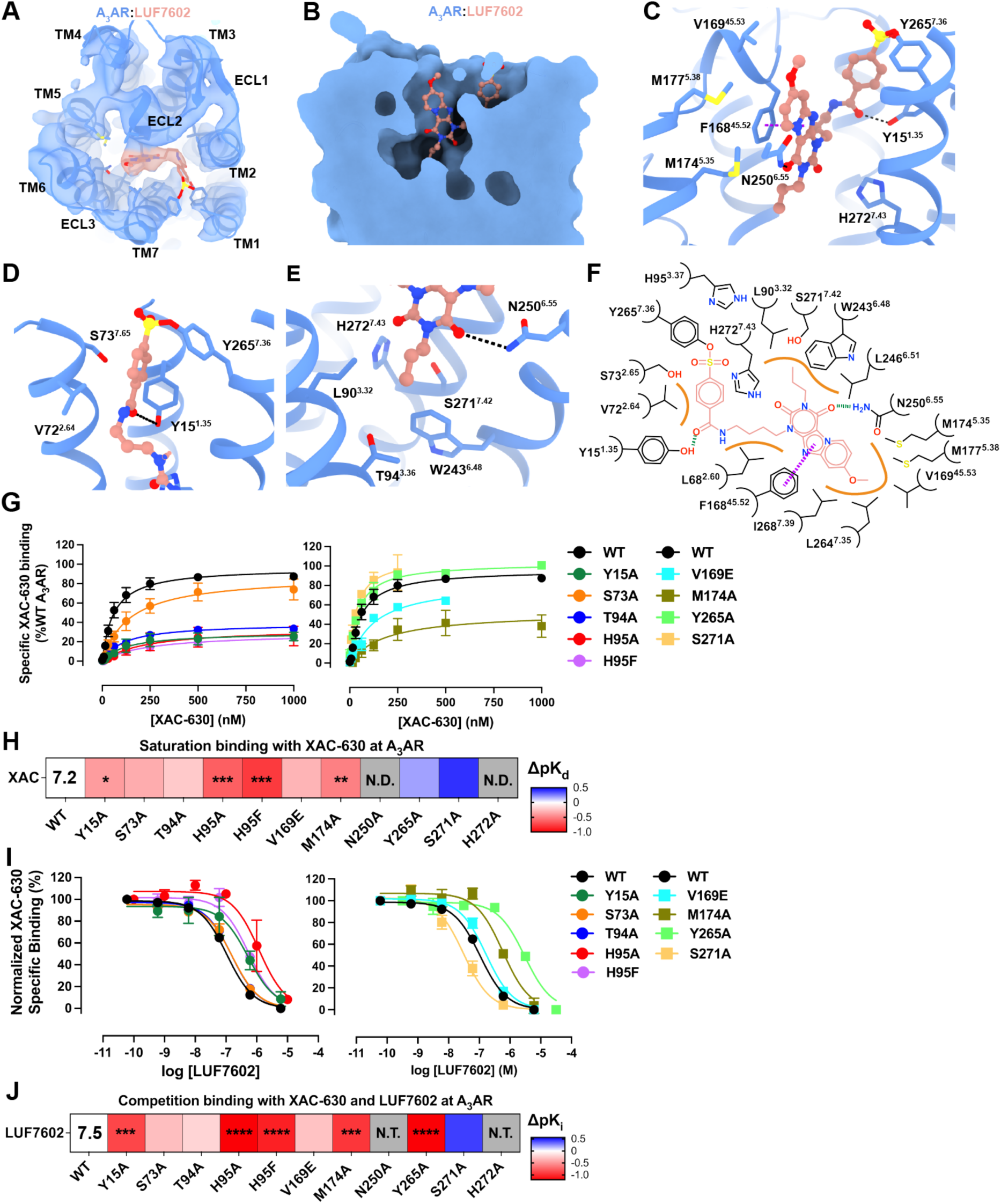
LUF7602 binding site at the A_3_AR. **(A)** Cryo-EM density map of A_3_AR (blue mesh) with bound LUF7602 (peach). **(B)** Surface representation of the A_3_AR orthosteric binding pocket with LUF7602 (peach sticks). **(C)** View of key residue interactions with LUF7602 in the binding pocket. **(D)** Detailed view of the covalent attachment of LUF7602 to Y265^7.36^ and hydrogen bond with _Y151.35._ **(E)** Hydrogen bond interactions between LUF7602 with N250^6.55^. **(F)** Schematic representation of LUF7602 interactions within the A_3_AR binding pocket. Hydrogen bonds are shown as green dashed lines, a π-π stacking interaction as purple dashed lines, and orange lines as hydrophobic interactions. **(G)** NanoBRET saturation specific binding curves for XAC-630 at wild-type A_3_AR and various mutants. Maximal specific binding was normalised to 100% of WT A_3_AR. Data shown are grouped mean ± SEM values. Grouped pK_d_ values and statistical analysis were from n=6 (WT), n=5 (Y15A, M174A), n=4 (H95A), and n=3 (S73A, T94A, H95F, V169E, N250A, Y265A, S271A, H272A). pK_d_ and Bmax values are reported in **Table 1**. **(H)** Heat map showing changes in binding affinity (ΔpK_d_) of XAC-630 to A_3_AR mutants relative to wild-type (WT) in saturation binding assays. N.D. = not determined due to no measurable response. The pK_d_ for XAC-630 at WT is overlaid on the heatmap. **(I)** Competitive binding curves showing the displacement of XAC-630 by LUF7602 at wild-type A_3_AR and selected mutants. The concentration of XAC-630 used in these experiments was approximate to the K_d_ determined in (G) with 200 nM used for all A_3_AR constructs except S271A where 40 nM was used. Grouped data are shown as mean ± SEM from n=3 experiments (n=4 for WT and Y15A) performed in duplicate. Grouped pK_i_ values and statistical analysis are shown in **Table 2**. **(J)** Heat map showing changes in binding affinity (ΔpK_i_) of LUF7602 competing with XAC-630 at A_3_AR mutants relative to WT. The pK_i_ for LUF7602 at WT is overlaid on the heatmap. Significant differences versus WT were determined by one-way ANOVA with a Dunnett’s multiple comparison test. P values: * = 0.01-0.05; ** = 0.001–0.01; *** = 0.0001-0.001; **** = < 0.0001. N.D.: Not determined. N.T.: Not tested. Blue indicates increased affinity, red indicates decreased affinity. Residue numbering follows Ballesteros-Weinstein convention.

**Table 1.**
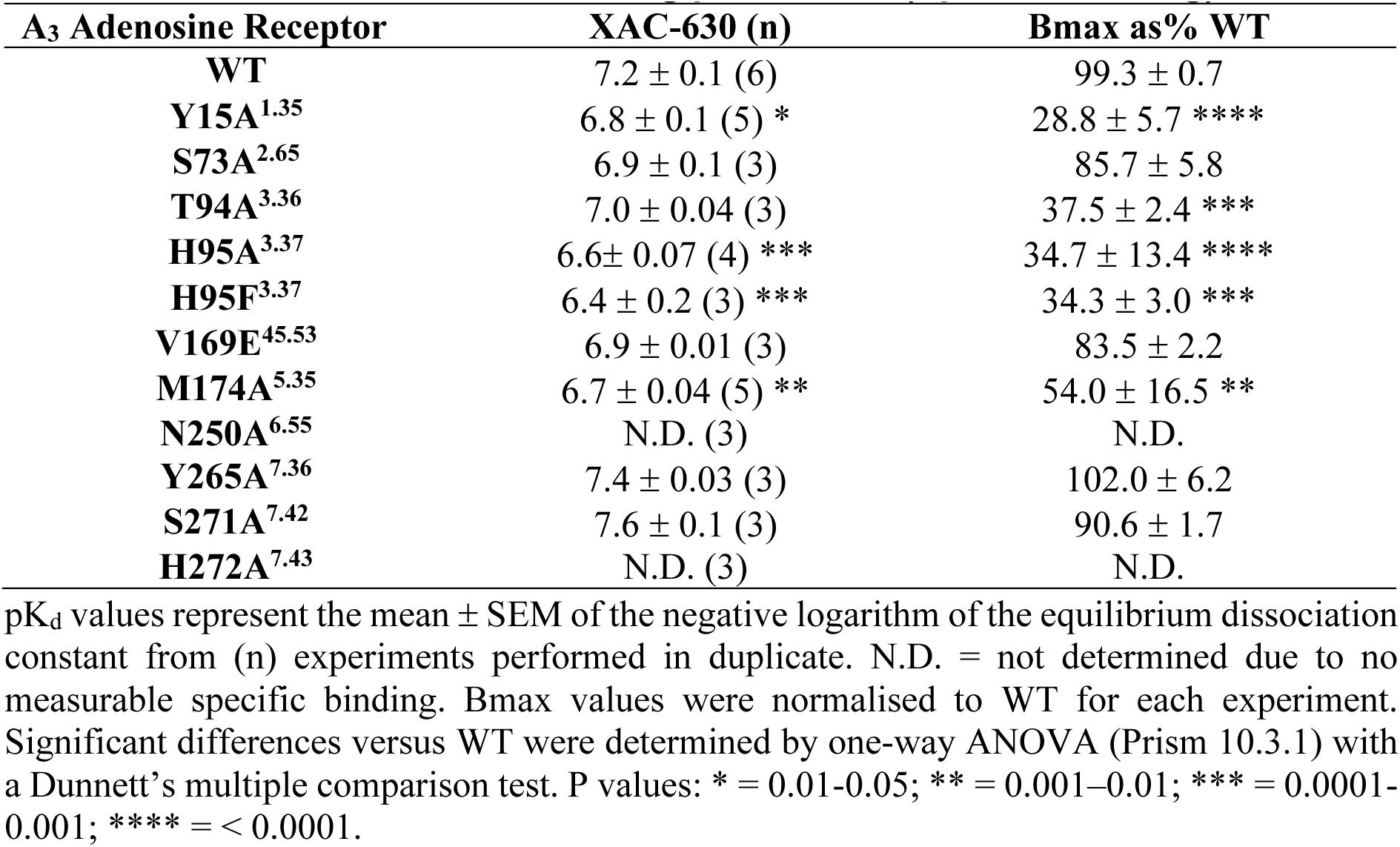
Nano-BRET saturation binding parameters (Specific Binding)

| **A<sub>3</sub> Adenosine Receptor** | **XAC-630 (n)** | **Bmax as% WT** |
| --- | --- | --- |
| **WT** | 7.2 ± 0.1 (6) | 99.3 ± 0.7 |
| **Y15A**<sup>1.35</sup> | 6.8 ± 0.1 (5) * | 28.8 ± 5.7 **** |
| **S73A**<sup>2.65</sup> | 6.9 ± 0.1 (3) | 85.7 ± 5.8 |
| **T94A**<sup>3.36</sup> | 7.0 ± 0.04 (3) | 37.5 ± 2.4 *** |
| **H95A**<sup>3.37</sup> | 6.6 ± 0.07 (4) *** | 34.7 ± 13.4 **** |
| **H95F**<sup>3.37</sup> | 6.4 ± 0.2 (3) *** | 34.3 ± 3.0 *** |
| **V169E**<sup>45.53</sup> | 6.9 ± 0.01 (3) | 83.5 ± 2.2 |
| **M174A**<sup>5.35</sup> | 6.7 ± 0.04 (5) ** | 54.0 ± 16.5 ** |
| **N250A**<sup>6.55</sup> | N.D. (3) | N.D. |
| **Y265A**<sup>7.36</sup> | 7.4 ± 0.03 (3) | 102.0 ± 6.2 |
| **S271A**<sup>7.42</sup> | 7.6 ± 0.1 (3) | 90.6 ± 1.7 |
| **H272A**<sup>7.43</sup> | N.D. (3) | N.D. |

pK<sub>d</sub> values represent the mean ± SEM of the negative logarithm of the equilibrium dissociation constant from (n) experiments performed in duplicate. N.D. = not determined due to no measurable specific binding. Bmax values were normalised to WT for each experiment. Significant differences versus WT were determined by one-way ANOVA (Prism 10.3.1) with a Dunnett's multiple comparison test. P values: \* = 0.01-0.05; \*\* = 0.001–0.01; \*\*\* = 0.0001-0.001; \*\*\*\* = < 0.0001.

**Table 2.** NanoBRET saturation binding parameters.

| **A<sub>3</sub> Adenosine Receptor** | **LUF7620**<br>**pK<sub>i</sub> (n)** | **Adenosine**<br>**pK<sub>i</sub> (n)** | **Piclidenoson**<br>**pK<sub>i</sub> (n)** | **NECA**<br>**pK<sub>i</sub> (n)** |
| --- | --- | --- | --- | --- |
| **WT** | 7.5 ± 0.05 (4) | 5.0 ± 0.10 (4) | 7.5 ± 0.08 (5) | 6.3 ± 0.1 (4) |
| **Y15A<sup>1.35</sup>** | 6.6 ± 0.2 (4) *** | 3.9 ± 0.1 (4) *** | 6.1 ± 0.2 (5) **** | N.D. (3) |
| **S73A<sup>2.65</sup>** | 7.2 ± 0.06 (3) | 4.60 ± 0.1 (3) | 6.6 ± 0.1 (4) ** | 5.4 ± 0.1 (3) * |
| **T94A<sup>3.36</sup>** | 7.3 ± 0.05 (3) | 4.9 ± 0.2 (3) | 7.1 ± 0.1 (4) | 4.6 ± 0.1 (3) *** |
| **H95A<sup>3.37</sup>** | 6.2 ± 0.2 (3) **** | 3.7 ± 0.05 (3) *** | 5.7 ± 0.3 (3) **** | 4.0 ± 0.1 (3) **** |
| **H95F<sup>3.37</sup>** | 6.4 ± 0.09 (3) **** | 3.9 ± 0.2 (3) *** | 5.7 ± 0.1 (4) **** | N.D. (4) |
| **V169E<sup>45.53</sup>** | 7.2 ± 0.05 (3) | 5.0 ± 0.08 (3) | 8.1 ± 0.2 (4) | 6.00 ± 0.2 (3) |
| **M174A<sup>5.35</sup>** | 6.5 ± 0.09 (3) *** | 4.3 ± 0.5 (3) | 7.9 ± 0.3 (4) | 5.3 ± 0.4 (3) ** |
| **Y265A<sup>7.36</sup>** | 6.3 ± 0.07 (3) **** | 4.50 ± 0.04 (3) | 6.7 ± 0.1 (4) **** | 5.0 ± 0.05 (3) *** |
| **S271A<sup>7.42</sup>** | 7.9 ± 0.09 (3) | N.D. (3) | 5.4 ± 0.1 (5) ** | N.D. (3) |

Values represent the mean ± SEM from (n) experiments performed in duplicate. N.D.= not determined due to no measurable specific binding. Significant differences versus WT were determined by one-way ANOVA (Prism 10.3.1) with a Dunnett's multiple comparison test. P values: \* = 0.01-0.05; \*\* = 0.001–0.01; \*\*\* = 0.0001-0.001; \*\*\*\* = < 0.0001.

We then assessed the ability of LUF7602 to compete with XAC-630 using a competition-binding NanoBRET assay (**Figure 2I,J**, **Table 2**). The Y265A^7.36^ mutant reduced LUF7602 affinity 15-fold, consistent with previously reported values (*39*). Nearby residue Y15^1.35^ forms a hydrogen bond with the amine group that links the xanthine core of LUF7602 to the reactive benzene-sulfonate warhead (**Figure 2D**). Loss of the hydrogen bond interaction via Y15A^1.35^ reduced LUF7602 affinity 10-fold. Poor EM density around the benzene-sulfonate group, Y265^7.36^, and Y15^1.35^ suggests conformational heterogeneity (**Figure 2A**), consistent with observations in the XAC-bound A_2A_AR structure (*33*). Residue N250^6.55^ is a conserved residue that forms hydrogen bonds with the heterocyclic rings of adenosine receptor agonists and antagonists. In the LUF7602-bound A_3_AR structure, N250^6.55^ forms a single hydrogen bond with the carbonyl-oxygen from the C^4^-position of the xanthine core (**Figure 2E,F**). We could not assess the impact of mutating N250^6.55^ due to a loss of XAC-630 binding, highlighting its general importance. The remaining interactions of LUF7602 with the A_3_AR were primarily hydrophobic interactions (**Figure 2F**). The H95A^3.37^ and H95F^3.37^ mutations disrupted LUF7602 binding ∼10-fold consistent with H95 being an important residue for the binding of agonists and antagonists (*64*). Similarly, the M174A^5.35^ mutant decreased LUF7602 affinity 10-fold (**Figure 2I,J**, **Table 2**). Finally, the affinity of LUF7602 for the S73A^2.65^ and T94A^3.36^ mutants were similar to WT A_3_AR, while the S271A^7.42^ caused a slight increase in affinity likely due to removal of a polar interaction and replacement with a hydrophobic interaction. Overall, these data support the binding mode of LUF7602 being due to diverse receptor-ligand interactions and provide opportunities to facilitate the design of higher affinity A_3_AR antagonists.

Comparison of xanthine-based antagonists bound to the orthosteric binding site of the A_1_AR, A_2A_AR, and A_3_AR revealed common and distinct binding interactions (**Figure 3**). Overall, the xanthine scaffolds occupy a similar pocket and form common interactions that include a π-π stacking interaction with conserved residue F^45.52^ and a hydrogen bond interaction with conserved residue N^6.55^. The orientation of the xanthine scaffolds was similar for all the ligands, with DU172 and PSB36 extending deeper into the orthosteric pocket (**Figure 3A**; red arrow). There was a slight tilt in the position of the xanthine scaffolds towards TM3 at each receptor subtype relative to the A_3_AR (**Figure 3A-C**; orange arrows). The largest tilt (∼42°) was observed with XAC at the A_2A_AR, followed by a 30° tilt for DU172 and PSB36 relative to the A_3_AR, while the other A_2A_AR xanthines were tilted 15-20°. Given that XAC was used as a fluorescent probe in our NanoBRET experiments, we performed induced fit docking with XAC and the A_3_AR structure (**Figure 3C**). The pose of the xanthine closely matched that of LUF7602, with the polar tail extending towards TM1, TM2, and TM7. However, the difference in the pose of XAC between the A_3_AR and A_2A_AR could be due to XAC binding in multiple conformations, as the electron density for XAC was not complete in the A_2A_AR structure (*33*).

**Fig. 3.**
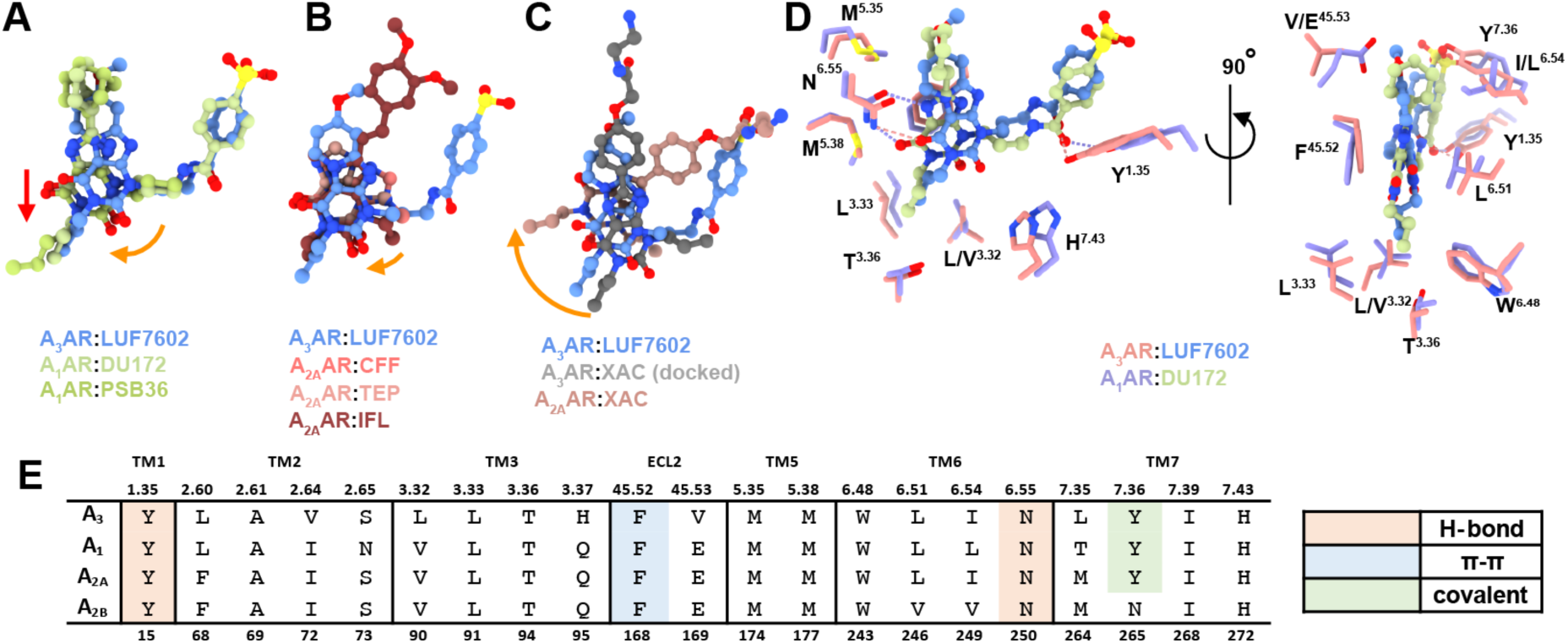
Comparison of the binding pose of adenosine receptor antagonists. **(A-C)** Binding poses of ligands from different antagonist-bound adenosine receptor structures. Red arrow indicates a displacement and orange arrows indicate rotations. (A) Comparison of A_3_AR:LUF7602 with A_1_AR:DU172 (PDB: 5UEN, light green) and A_1_AR:PSB36 (PDB: 5N2S, light green) (B) Comparison of A_3_AR:LUF7602 with A_2A_AR:CFF (PDB: 5MZP, peach), A_2A_AR:TEP (PDB: 5MZJ, pink), A_2A_AR:IFL (PDB: 8GNG, red) (C) Comparison of A_3_AR:LUF7602 with A_3_AR:XAC (docked) and A_2A_AR:XAC (PDB: 3REY, dark pink). **(D)** Detailed view of ligand-receptor interactions for A_3A_R:LUF7602 and A_1_AR:DU172, showing key residues involved in binding. **(E)** Detailed view of ligand-receptor interactions for A_3_AR:LUF7602 and A_1_AR:DU172, showing key residues involved in binding.

DU172 and LUF7602 are modestly selective A_1_AR and A_3_AR covalent antagonists with similar chemical structures, binding affinities (DU172: pK_i_ = 7.4 at A_1_AR; LUF7602: pK_i_ = 7.2 at A_3_AR), and binding modes (**Figure 3D**). Both ligands form a covalent linkage with residue Y^7.36^ and have a propyl group on N^3^ that forms hydrophobic contacts with L^3.33^, T^3.36^, L^6.51^, M177^5.38^, and W^6.48^. In addition, both ligands have chemical groups that extend off the C^8^ position into a pocket created by ECL2, TM6, and TM7. In the case of DU172, the larger piperazine forms hydrophobic interactions with E172^45.53^, M177^5.35^, L253^6.54^, N254^6.55^, T257^6.58^, and T270^7.35^. This pocket is less conserved among adenosine receptor subtypes, and residue T270^7.35^ was shown to contribute to the subtype selectivity of DU172. In contrast, the methoxy group of LUF7602 only interacts with V169^45.53^ and L264^7.35^. Residue V169^45.53^ is an E at all other adenosine receptor subtypes, which we hypothesised would cause a steric clash with LUF7602 (**Figure 3D**). However, the V169E^45.53^ mutation did not affect LUF7602 binding (**Figure 2I,J**), suggesting the residue adopted a different rotamer.

### Active-state structures of the A_3_AR

Next, we sought to determine the active-state structures of the A_3_AR in complex with G protein. To determine the structure of the A_3_AR bound to endogenous agonist adenosine, we used the single-chain antibody scFv16 (*65*) and the dominant negative form of Gα_i1_ (DNGα_i1_) to stabilise the complex (*66, 67*). Single-particle cryo-EM analysis of the purified complex samples resulted in a 2.9 Å map providing detailed insights into the A_3_AR-DNG_i1_-scFv16-adenosine complex (**Figure 4A,B, S5B, S6, Table S1)**.

**Fig. 4.**
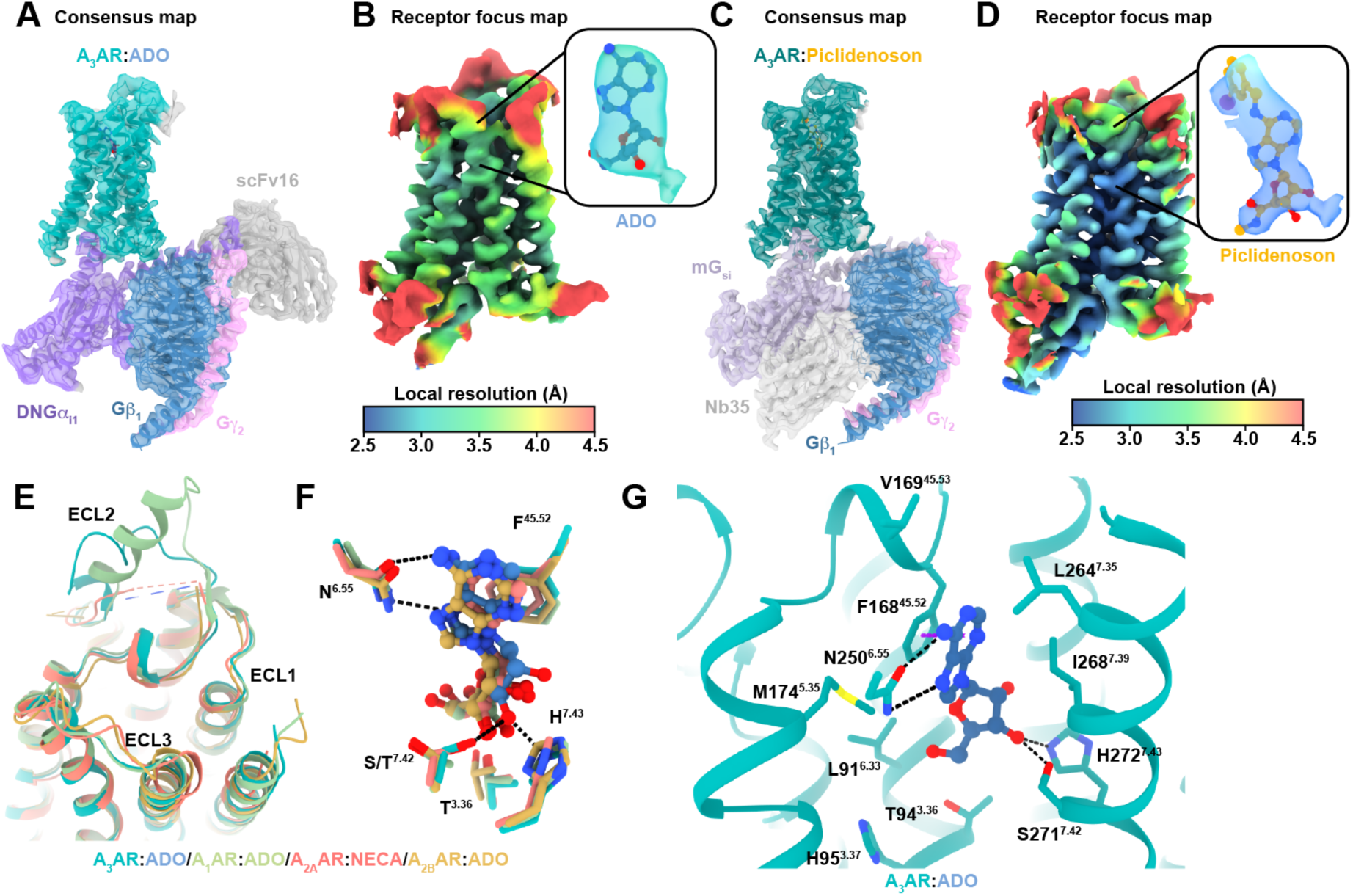
Comparison of cryo-EM structures in the active conformation. **(A)** Cryo-EM density map and atomic model of the active A_3_AR:adenosine (ADO) complex, showing the receptor (green), ADO (blue), DNGα_i1_ (purple), Gβ_1_ (dark blue), Gγ_2_ (pink), and scFv16 (grey). **(B)** Local resolution receptor focused cryo-EM map with inset of the density around ADO. **(C)** Cryo-EM density map and atomic model of the active A_3_AR:Piclidenoson complex, showing the receptor (dark green), Piclidenoson (orange), mG_si_ (light purple), Gβ_1_ (dark blue), Gγ_2_ (pink), and Nb35 (grey). **(D)** Local resolution receptor focused cryo-EM map with inset of the density around Piclidenoson. **(E).** Extracellular view of aligned structures of A_3_AR:ADO (cyan), A_1_AR:ADO (PDB: 7LD4, green), A_2A_AR:NECA (PDB: 6GDG, red), and A_2B_AR:ADO (PDB: 8HDP, orange). **(F)** Comparison of adenosine binding in A_3_AR:ADO (cyan), A_1_AR:ADO (PDB: 7LD4, green), A_2A_AR:ADO (PDB: 2YDO, red), and A_2B_AR:ADO (PDB: 8HDP, orange), highlighting key interacting residues. **(G)** Detailed view of A_3_AR:ADO interactions, showing key residues involved in ligand binding.

To obtain a higher-resolution structure with Piclidenoson-bound to the A_3_AR, we used an engineered mini-G_si_ construct (*68*) that was stabilized using nanobody Nb35 (*65, 69*) **(Figure S7A)**. For the A_3_AR-mG_si-_Nb35-Piclidenoson complex, we achieved a global resolution of 2.5 Å. Following local refinement, the receptor map reached a resolution of 3.3 Å (**Figure 4C,D, S5C, S7, Table S1)**. Both the ECL and ICL regions were well-resolved and displayed nearly identical conformations, allowing clear identification of most receptor residues except for residues 208-225 in ICL3, which remained disordered.

### Comparison of the active conformation at adenosine receptors

The adenosine- and Piclidenoson-bound A_3_AR structures closely resemble other active-state adenosine receptor structures with RMSDs of less than 1.0 Å. The TM helices aligned well, with minor divergence at the top of TM7 in the A_2A_AR and A_2B_AR due to a longer ECL3 (**Figures 4E**). Despite moderate overall sequence similarity, the pose and position of adenosine in the orthosteric site were highly similar across all adenosine receptor structures (**Figure 4F**). The ribose ring extends deeply into the binding site, anchored by a hydrogen bond network involving residues in TM 3, 6, 7 and ECL2. The conserved interaction network between adenosine and ARs includes a π-stacking interaction with F^45.52^, a bidentate hydrogen bond with N^6.55^, hydrogen bonds between the 3’, and 5’ hydroxyl groups with S/T^7.42^ and H^7.43^, and numerous hydrophobic interactions (**Figure 4F,G**).

Piclidenson, an A_3_AR-selective agonist, is a larger molecule than adenosine and extends from the orthosteric site out towards ECL2 and TMs4-6. Piclidenson differs from adenosine by two key substituents: a methylcarboxamide group extending from the C5’ position and an N^6^-iodobenzyl group (**Figure 5A-C**) (*70, 71*). The methylcarboxamide group forms a hydrogen bond with T94^3.36^, with the methyl group extending into a hydrophobic pocket composed of residues L91^3.33^, H95^3.37^, S181^5.42^, I186^5.47^, and W243^6.48^ (**Figure 5D**). Non-conserved residues H95^3.37^ and S181^5.4^ restrict the size of the orthosteric pocket. The importance of this pocket is underscored by the 30-fold decrease in 2-Cl-Piclidenoson (i.e., Namodenoson) binding at the H95^3.37^ alanine mutation (*72*). The pocket’s proximity to the rotamer toggle switch residue W243^6.48^ suggests a role in receptor activation. The N^6^-iodobenzyl group occupies a hydrophobic pocket formed by TMs 5-7 and ECL2, making various hydrophobic interactions (**Figure 5E**). Notably, the iodine atom points towards the backbone carbonyl of M172^45.56^ in an orientation (θ_1_ = 126°, θ_2_ = 103°) and distance (3.4 Å) that favours a halogen bond interaction (*73*). Structure-activity relationships support this halogen bond, as replacement of the iodine atom with H or Cl decreased binding affinity by16- and 20-fold, respectively (*71*).

**Fig. 5.**
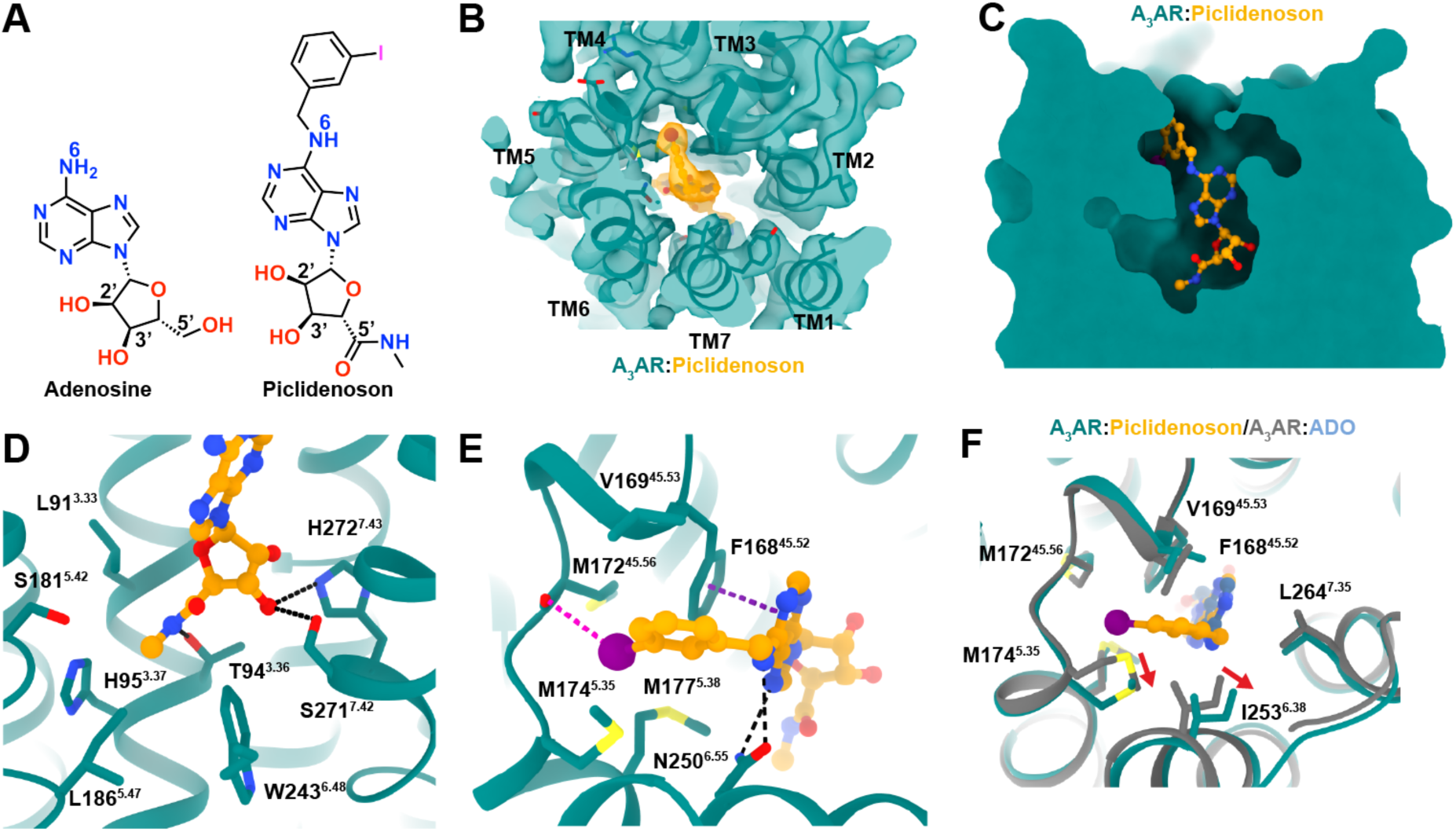
The Piclidenoson binding site. **(A)** Chemical structures of adenosine and Piclidenoson with key atoms numbered. **(B)** Cryo-EM map of A_3_AR Piclidenoson**-**bound binding site. **(C)** Surface representation of the A_3_AR orthosteric binding pocket with Piclidenoson shown as orange sticks. **(D-E)** Detailed view of A_3_AR:Piclidenoson interactions, showing key residues involved in ligand binding near (D) the ribose group and (E) the N^6^-iodobenzyl group. **(F)** Comparison of the position of residue M174^5.35^ in the adenosine-bound (coloured gray) and Piclidenoson-bound structures.

To validate the observed agonist binding modes, we measured the binding affinity of adenosine and Piclidenoson using a competition NanoBRET binding assay with XAC-630. We also tested NECA because it is chemically similar to adenosine and Piclidenoson with an ethylcarboxamido group that extends from the C5’ position (**Figure 6A-C**, **Table 2**). The agonists had a binding affinity rank order of Piclidenoson > NECA > adenosine, consistent with previous studies and with Piclidenoson’s more extensive interactions with the receptor compared to adenosine (**Figure 6D,E**). Residue S271A^7.42^ caused the largest loss of binding across all three agonists due to the loss of interaction with the 3’ OH group. Similarly, H95A^3.37^ and H95F^3.37^ significantly affected binding of all three agonists, particularly NECA, suggesting these mutations impact the orthosteric site. Interestingly, we observed that Y15A^1.35^ significantly affected the binding for all three agonists despite not forming any direct interactions. This residue forms part of a hydrogen bond network between TM1, TM2, and TM7 that helps coordinate H272^7.43^ with the ribose group (**Figure 7E**). Mutations S73A^2.65^ and Y265A^7.36^ had similar effects, though less apparent with adenosine. Residue Y265^7.36^ contributes to the hydrogen bond network by forming a π-stacking interaction with Y15A^1.35^. The role of S73^2.65^ was more subtle, but it can hydrogen bond to Y265^7.36^ when adopting a different rotamer (**Figure 7E**).

**Fig. 6.**
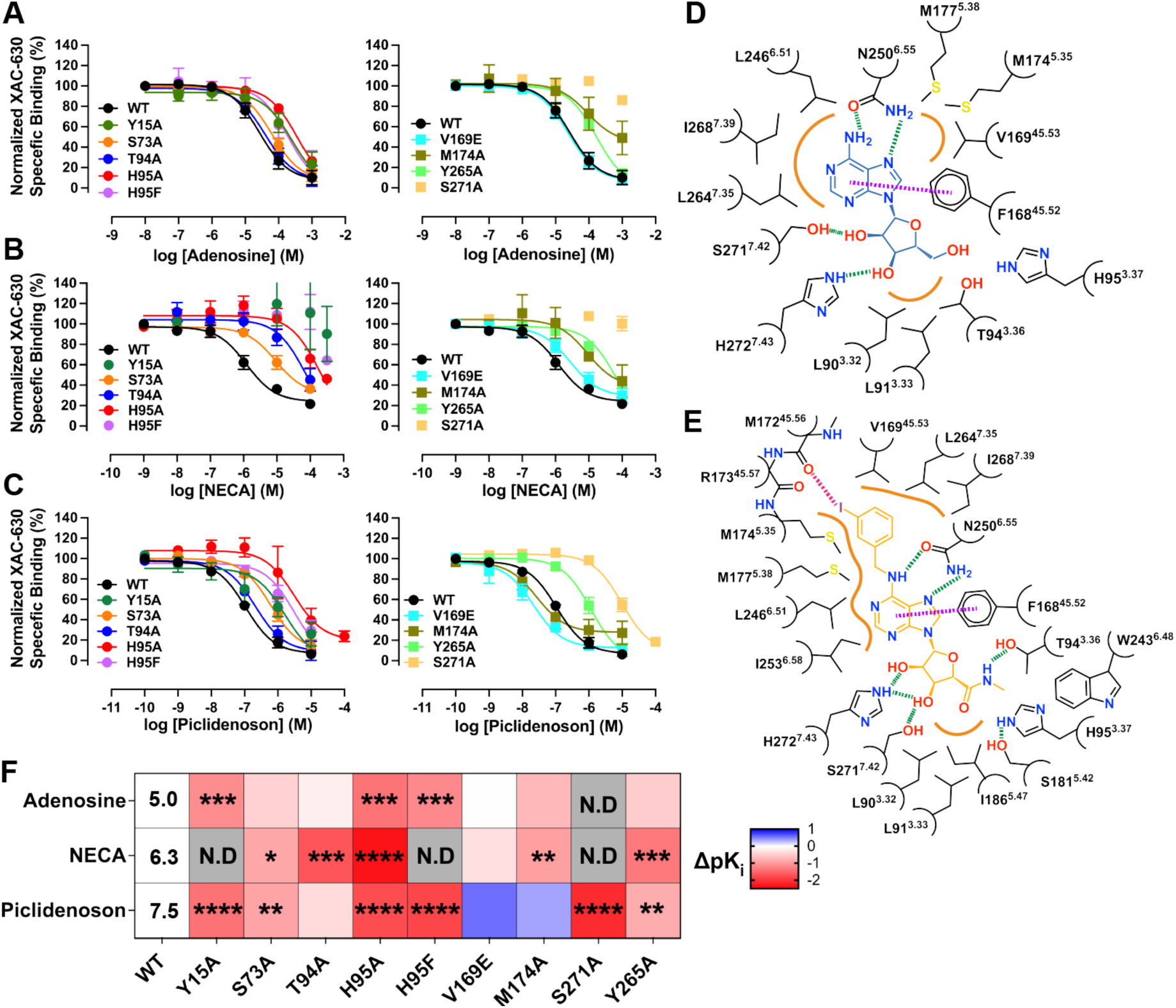
Key residues in the agonist-binding site. **(A-C)** Competitive binding curves showing the displacement of XAC-630 by (A) adenosine, (B) NECA, and (C) Piclidenoson at wild-type (WT) and mutant A_3_AR constructs. Grouped data are shown as mean ± SEM experiments performed in duplicate. Grouped pK_i_ values, experimental numbers, and statistical analysis are shown in **Table 2**. **(D, E)** 2D interaction diagram of (D) the A_3_AR:ADO complex and (E) the A_3_AR:Piclidenoson complex. Hydrogen bonds are shown as green dashed lines, a π-π stacking interaction as purple dashed lines, halogen bond as a pinked dashed line, and orange lines as hydrophobic interactions. Residue numbering follows Ballesteros-Weinstein convention. **(F)** Heat map showing the effects of various A_3_AR mutations on binding affinity (ΔpK_i_) for adenosine, NECA, and Piclidenoson. The pK_i_ for adenosine, NECA, and Piclidenoson at WT is overlaid on the heatmap. Significant differences versus WT were determined by one-way ANOVA with a Dunnett’s multiple comparison test. P values: * = 0.01-0.05; ** = 0.001–0.01; *** = 0.0001-0.001; **** = < 0.0001. N.D.: Not determined. Blue indicates increased affinity, red indicates decreased affinity.

**Fig. 7.**
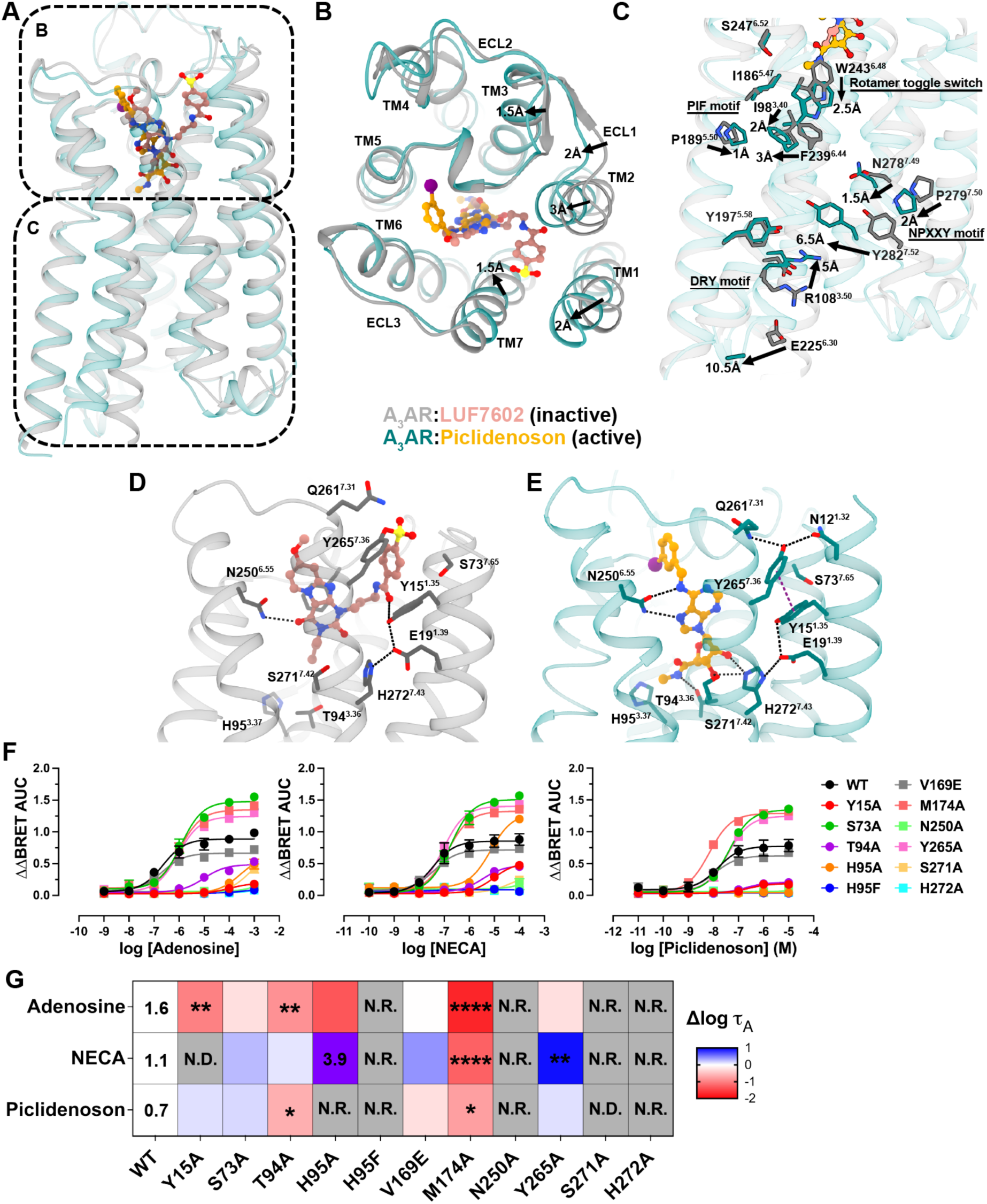
Activation mechanism of the A_3_AR. **(A)**. Overall structure of A_3_AR in active (green) and inactive (light blue) states. Top panel shows the extracellular view in (B), and bottom panel shows the side view in (C). **(B)**. Detailed comparison of active and inactive A_3_AR structures, highlighting key conformational changes viewed from the extracellular surface. Arrows indicate the direction and magnitude of movements. **(C)**. Close-up view of the intracellular region, showing key residues and motifs involved in receptor activation. Signalling motifs such as the PIF motif, rotamer toggle switch, NPXXY motif, and DRY motif are labeled. Arrows indicate conformational changes with distances. **(D,E)** The binding pocket of (D) the inactive A_3_AR bound to LUF7602 and (E) the active A_3_AR bound to Piclidenoson. Key residues involved in ligand interactions are shown and labeled. Hydrogen bonds are depicted as black dashes and π-π stacking interactions as purple dashed lines. **(F)** Gα_i1_ activation using the TruPath assay. Trupath experimental data for each agonist was baseline-corrected to the initial baseline and then to the buffer control (ΔΔ) followed by calculating the area under the curve (AUC) for each response (ΔΔBRET AUC). Data points represent grouped mean ± SEM values. Data shown were fit against a three-parameter log[agonist] vs. response model. **(G)** Heat map showing the effects of various A_3_AR mutations on signalling efficacy (Δlog 1_A_) for adenosine, NECA, and Piclidenoson. The log 1_A_ for adenosine, NECA, and Piclidenoson at WT and NECA at H95A are overlaid on the heatmap. Significant differences versus WT were determined by one-way ANOVA with a Dunnett’s multiple comparison test. P values: * = 0.01-0.05; ** = 0.001–0.01; *** = 0.0001-0.001; **** = < 0.0001. N.D.: 1_A_ not determined due to no lack of a K_A_ value. N.R.: No response. Blue indicates increased efficacy, red indicates decreased efficacy, purple square for NECA indicates a much higher log 1_A_ value. Values are reported in **Table 3**.

**Table 3.** Gα_i1_ activation assay (TruPath)

| **A<sub>3</sub> Adenosine Receptor** | **Adenosine**<br>**log $\tau$ (n)** | **Piclidenoson**<br>**log $\tau$ (n)** | **NECA**<br>**log $\tau$ (n)** |
| --- | --- | --- | --- |
| **WT** | 1.6 ± 0.2 (3) | 0.7 ± 0.1 (3) | 1.1 ± 0.2 (3) |
| **Y15A**<sup>1.35</sup> | 0.6 ± 0.3 (3) ** | 0.8 ± 0.3 (3) | N.D. (3) |
| **S73A**<sup>2.65</sup> | 1.4 ± 0.07 (3) | 0.8 ± 0.07 (3) | 1.4 ± 0.07 (3) |
| **T94A**<sup>3.36</sup> | 0.8 ± 0.09 (3) * | 0.01 ± 0.07 (3) * | 1.2 ± 0.1 (3) |
| **H95A**<sup>3.37</sup> | 0.3 ± 0.3 (3) *** | N.D. (3) | 3.9 ± 0.02 (3) **** |
| **H95F**<sup>3.37</sup> | N.R. (3) | N.R. (3) | N.R. (3) |
| **V169E**<sup>45.53</sup> | 1.6 ± 0.1 (3) | 0.4 ± 0.09 (3) | 1.6 ± 0.1 (3) |
| **M174A**<sup>5.35</sup> | -0.1 ± 0.2 (3) **** | -0.1 ± 0.2 (3) * | -0.1 ± 0.2 (3) *** |
| **N250A**<sup>6.55</sup> | N.R. | N.R. | N.R. (3) |
| **Y265A**<sup>7.36</sup> | 1.4 ± 0.07 (3) | 0.8 ± 0.06 (3) | 2.1 ± 0.07 (3) ** |
| **S271A**<sup>7.42</sup> | N.D. (3) | N.R. (3) | N.R. (3) |
| **H272A**<sup>7.43</sup> | N.R. (3) | N.R. (3) | N.R. (3) |

In contrast the T94A^3.36^ mutation only significantly reduced the binding of NECA, suggesting it forms a key interaction with the ethylcarboxamido group (**Figure 6F**). Similar to LUF7602, the ECL2 V169E^45.53^ mutation was designed to also test the selectivity of Piclidenson. However, the mutation resulted in a 10-fold increase in binding affinity for Piclidenoson, suggesting a less clear role (**Figure 6F**).

Mutation of residue M174A^5.35^ had paradoxical effects. With adenosine and NECA, M174A^5.35^ slightly reduced affinity and showed non-competitive displacement (**Fig 6A,B**). For Piclidenoson, M174A^5.35^ increased affinity and enhanced displacement of XAC-630 (**Fig 6C**). This result aligns with our cryo-EM data, which shows that the conformation of M174^5.35^ is influenced by the specific ligand occupying the orthosteric binding site (**Fig 5F).** In the Piclidenoson-bound structure M174^5.35^ is pushed back towards TM6 by ∼2-3Å compared to adenosine- and LUF7602-bound structures and to M^5.35^ in structures of the A_1_AR, A_2A_AR, and A_2B_AR. This suggests that M174^5.35^ functions as a gatekeeper residue, controlling access to a cryptic extracellular pocket. This cryptic pocket may also facilitate non-specific binding of XAC-630 as the position of the 2-aminoethyl-acetamide group of XAC appeared flexible in our docking of to the A_3_AR. This could potentially explain the observed non-competitive inhibition with the M174A^5.35^ mutation. Interestingly, in a recent structure Piclidenoson -bound A_3_AR structure (PDB: 8X16), the N^6^-iodobenzene group was modelled oriented towards the solvent **(Figure S8),** with no corresponding changes to the conformation of M174^5.35^. However, poor map density in the ECL region of 8X16 suggests potential uncertainties in the side chain and ligand modelling.

### Activation mechanism

The comparison of the inactive, LUF7602-bound, and active, agonist-bound A_3_AR complex structures offers insight into the mechanism of A_3_AR activation (**Figure 7A-C**). Adenosine and Piclidenoson bind deeper in the orthosteric site than LUF7602, triggering conformational changes characteristic of class A GPCRs (*74*). These changes involve a 2.5 Å downward shift in the rotamer toggle switch residue W243^6.48^, leading to signal propagation through the PIF, NPxxY, and DRY motifs, followed by an ∼11 Å outward movement of TM6 that is typical of class A G_i_ coupled receptors (*75*). While LUF7602, adenosine, and Piclidenoson interact similarly with orthosteric site residues (F168^45.42^, N250^6.55^, H272^7.43^), their binding modes and resulting A_3_AR conformations differ significantly. In the active state, TM1, TM2, ECL1, and TM7 move inward at the extracellular side (**Figure 7B**), breaking the ECL2 anti-parallel β-sheet observed in the inactive conformation.

A key feature of the active conformation is an extensive hydrogen bond network extending from the top of TM7 through TM1, TM2, and down to the ribose binding site (**Figure 5D,E**). Specifically, Q261^7.31^ and N12^1.32^ form hydrogen bonds with Y265^7.36^, positioning it to form a π-π stacking interaction with Y15^1.35^. Residue Y15^1.35^ forms a hydrogen bond with E19^1.39^, which forms a hydrogen bond with H272^7.43^ that positions H272^7.43^ for interaction with the 2’ and 3’ OH groups of adenosine and Piclidenoson. The 3’OH group of the agonists also interact with S271^7.42^, while the 5’ group may interact with T94^3.3.6^ (**Figure 7E**). We note that disruption of this hydrogen bond network by mutation of Y15^1.35^, S73^2.65^, S271^7.42^, and Y265^7.36^ significantly reduced agonist binding (**Figure 6F**). Notably, this extensive hydrogen bond network is absent in the LUF7602-bound structure, partly due to LUF7602’s covalent interaction with Y265^7.36^ disrupting the conformation of nearby residues (**Figure 7D**). Consequently, LUF7602 forms fewer interactions with the receptor compared to adenosine and Piclidenoson (**Figure 2F vs 6D,E)**. The uniqueness of this extensive hydrogen bond network to A_3_AR, not observed in other AR subtypes, suggests a distinct activation mechanism for this receptor subtype.

To test the role of residues surrounding the agonist-binding site on receptor activation, we used the BRET-based Trupath assay (*76*) to measure the proximal activation of Gα_i1_ (**Figure 7F**). The efficacy of the agonists (1_A_) were calculated using the Black–Leff operational model of agonism (*77*) and corrected for differences in receptor expression (*78*) (**Figure 7G**, **Table 3**). The agonists had a rank order of efficacy with adenosine > NECA > Piclidenoson consistent with prior observations that higher affinity agonists may have lower signalling efficacy (*79*). Similar to our findings from binding experiments, the N250A^6.55^ and H272A^7.43^ mutants did not signal suggesting these mutations impair receptor function. Similarly, NECA and Piclidenoson could not activate S271A^7.42^, while adenosine produced a weak response that could not be quantified due to no measurable binding affinity (**Figure 6A).** In addition, H95F^3.37^ did not produce a measurable response suggesting that although agonists could bind to this receptor mutant, they could not activate the receptor. The H95A^3.37^ mutation, however displayed ligand-dependent effects producing no response for Piclidenoson, a small response for adenosine, and a robust response for NECA despite a 100-fold loss in binding affinity. The increase in efficacy for NECA at H95A^3.37^ is likely related to its juxtaposition to the ethylcarboxamido group and the rotamer toggle switch residue W243^6.48^. Similarly, there was a ligand-dependent increase in the efficacy of NECA at the Y265A^7.36^ suggesting that Y265A^7.36^ reduces agonist binding but does not alter signalling. Nearby mutation Y15A^1.35^ reduced adenosine efficacy to a similar level as Piclidenoson indicating a small effect on signalling. Finally, the M174A^5.35^ mutation significantly reduced agonist efficacy to the same levels (**Table 3**) indicating M174^5.35^ is important for the binding and signalling of agonists, but with less of an effect on Piclidenoson. Overall, these orthosteric site mutations reveal distinct roles for binding site residues in receptor activation, with some mutations completely abolishing signaling while others show ligand-dependent effects on efficacy.

### G protein interface

Adenosine receptor subtypes exhibit different G protein coupling preferences (*80*), with A_1_AR and A_3_AR preferentially coupling to G_i_ proteins and A_2A/2B_AR coupling to G_s_ proteins (**Figure 8A**). Our study employed two different G protein constructs: a dominant negative G_i1_ for the adenosine-bound structure **(Figure S2A)** and a mini-G_si_ chimera for the Piclidenoson-bound structure **(Figure S7A)**. Comparing the adenosine- and Piclidenoson-bound structures revealed a similar set of interactions with the last five residues of the G protein α5-helix and the A_3_AR core (**Figure 8B-D**). Beyond these five residues, the interactions between G proteins and A_3_AR diverge, with the α5-helix of G_i1_ rotating away by 5Å near the end of the α5-helix (**Figure 8E**). Globally, this can be viewed as a rotation between the G proteins around the core of the receptor that results in a large displacement between the αN-helices (**Figure 8F**). On the one hand, these differences may be due to the different G protein constructs that were used. For example, the conformation of the α5-helix in the adenosine-A_3_AR-G_i1_ structure was more similar to the A_1_AR-G_i1_ structure and a recent Piclidenoson-A_3_AR-G_i_ structure (PDB: 8X16), while the α5-helix in our Piclidenoson-A_3_AR-mini-G_si_ structure was more similar to the A_2A/2B_AR-G_s_ structures. On the other hand, the conformation of the αN-helix in the adenosine-A_3_AR-G_i1_ appears to be an outlier compared to the other adenosine receptor structures, suggesting this could be specific to adenosine-A_3_AR-G_i1_. Despite the two Piclidenoson-bound structures using different G protein constructs and stabilizing methods, Nb35 versus LgBit-HiBit tethering (*81*), a similar number of contacts were made between the A_3_AR and G protein (**Figure 8C**). Notably, the adenosine-bound structure shows fewer receptor-G protein contacts compared to the Piclidenoson-bound structure, potentially revealing ligand-dependent A_3_AR-G protein conformations. We caveat these statements by highlighting the differences in G protein constructs and stabilizing methodologies that are commonly used in cryo-EM studies, and further studies will be required.

**Fig. 8.**
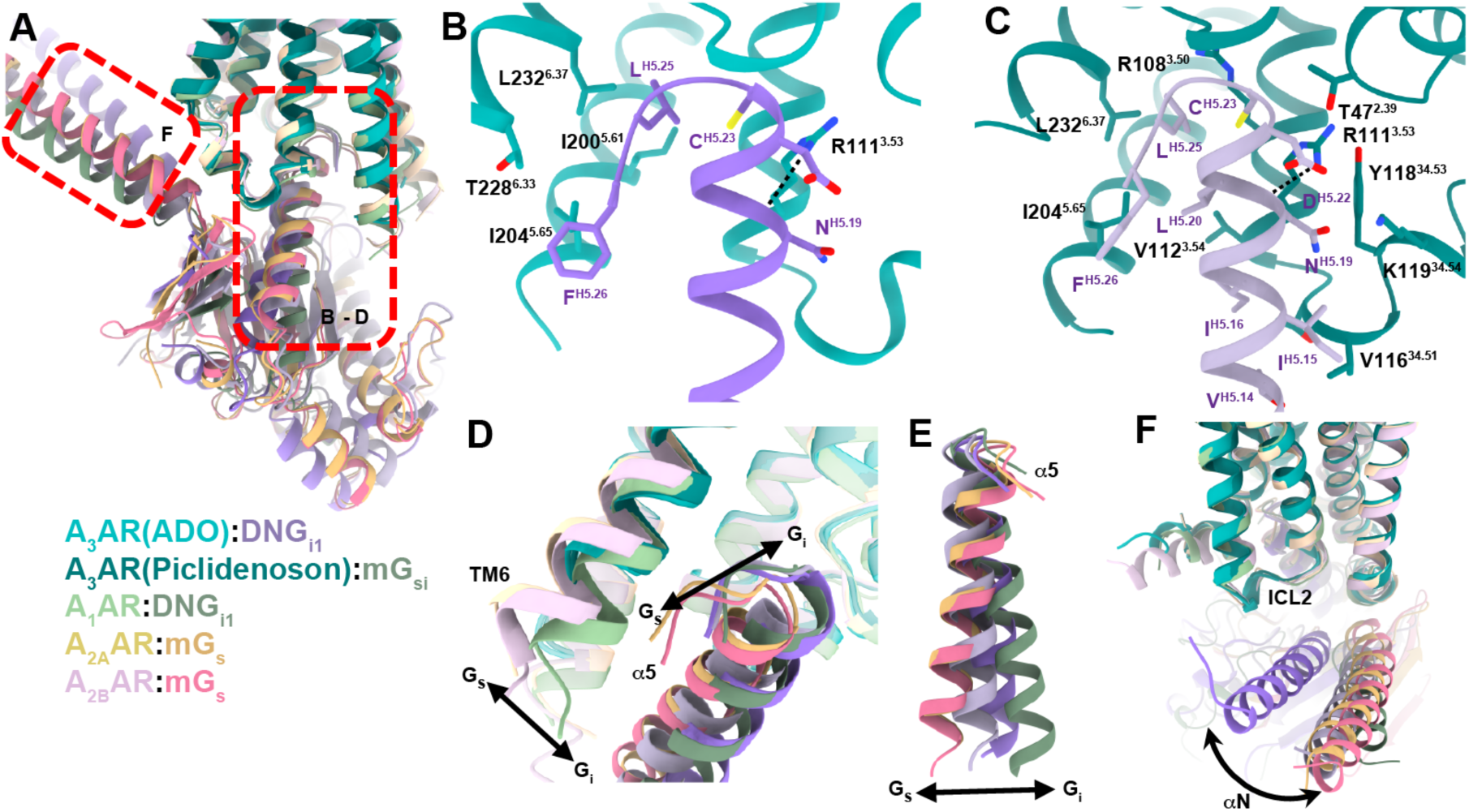
Comparison of the adenosine receptor G protein interface. **(A)**. Comparison of adenosine receptor – Gα complexes. View is of the interaction of Gα with the receptor. Proteins are coloured according to the legend. Red dashed box indicates regions shown in panels B-F. Structures shown and coloured as: A_3_AR:ADO:DNG_i1_ (coloured cyan and purple, PDB: 9EBH); A_3_AR:Piclidenoson:mG_si_ (coloured green and light purple, PDB: 9EBI); A_1_AR:ADO:DNG_i1_ (coloured light green and dark green, PDB: 7LD4); A_2A_AR:NECA:mG_s_ (coloured yellow and orange, PDB: 6GDG); and A_2B_AR:ADO:mGs (coloured light pink and red, PDB: 8HDP, orange). **(B,C)**. Detailed view of the A_3_AR:G protein interface from (B) the adenosine-bound A_3_AR-DNG_i1_ complex, and (C) the Piclidenoson-bound A_3_AR-mGsi complex. Receptor residues are shown shades of green, and G protein residues in shades of purple. Receptor residues are numbered according to Ballesteros and Weinstein numbering and G proteins are numbered according to the CGN scheme. **(D)**. Comparison of G protein coupling in different adenosine receptor structures, highlighting the positions of TM6 and the α5 helix. **(E)**. Overall Comparison of the α5 helix orientation in different G_s_ and G_i_ protein bound adenosine receptor structures. (F). Comparison of the αN helix from the adenosine-bound A_3_AR structure (purple) with other adenosine receptor:G protein complexes.

## Discussion

The human A_3_AR has attracted considerable interest as a drug target for treating various indications, as evidenced by multiple drugs now progressing through clinical trials. Given the clinical potential of the A_3_AR significant effort was invested into identifying potent and selective A_3_AR agonists, antagonists, and allosteric modulators (*17, 82–85*). To understand how potential drugs bind and modulate the activity of the A_3_AR we determined cryo-EM structures of the A_3_AR in the inactive and active conformations.

The structure of A_3_AR bound to the covalent antagonist LUF7602 revealed a binding mode that shares similarities with other xanthine-based antagonists bound to A_1_AR and A_2A_AR including the A_1_AR covalent antagonist DU172 (*32, 33*). The covalent attachment to Y265^7.36^, while crucial for structure determination, appears to disrupt optimal interactions with key residues N250^6.55^ and H272^7.43^. Although the covalent properties of DU172 and LUF7602 were essential to structure determination, both ligands have lower binding affinities than their parent molecules, DPCPX and a pyrido-[2,1-f]purine-2,4-dione (*40, 86*). This suggests that the covalent attachment of the ligands may result in less favourable interactions with the receptor. Future structures or computational studies with the parent molecules could provide further insight and aid the discovery of better antagonists.

We were not successful in determining a cryo-EM structure of the A_3_AR bound to the antagonist MRS1220. This was potentially a limitation of using the BRIL fusion approach (*43*) where an antibody fragment and Nb provide stability to the complex. Though these regions align well, there is a loss in resolution moving towards the more dynamic regions of the A_3_AR, which include the orthosteric site and the ECL region. In contrast, the addition of the covalent antagonist LUF7602 provided sufficient stability for structure determination, albeit the overall resolution surrounding the ligand binding site was at moderate resolution. Other methods to stabilise the inactive state of the receptor, while providing a fiducial marker such as Nb6 (*87*) or fusions with the C-terminus (*88*), may provide alternative approaches for future efforts.

The structures of A_3_AR bound to the endogenous agonist adenosine and the clinically relevant agonist Piclidenoson provide valuable insights into the molecular basis of agonist recognition and selectivity. The binding mode of adenosine is highly conserved across AR subtypes, consistent with its role as the endogenous ligand. In contrast, Piclidenoson’s unique substituents, particularly the N^6^-iodobenzyl group, exploit subtype-specific features of the A_3_AR binding pocket. Comparison of our Piclidenoson-bound structure to other A_3_AR and adenosine receptor structures revealed that the N^6^-iodobenzyl group bound to a cryptic pocket due to the movement of a conserved gatekeeper residue M174^5.35^. Cryptic pockets are binding sites that are rarely observed in experimental structures, because they are formed due to protein fluctuations that create the cryptic pocket, in this case M174^5.35^. Instead, these pockets are typically observed via molecular dynamic simulations (*89–93*). The discovery of a cryptic pocket accommodating the N^6^-iodobenzyl group, offers an exciting opportunity for design of more highly selective A_3_AR agonists, but also underscores the value of determining multiple cryo-EM structures of the same protein-ligand complex. This approach can reveal dynamic features and conformational heterogeneity that might be missed in a single structure, highlighting the utility of comprehensive structural analyses in GPCR research.

Comparison of our inactive and active A_3_AR structures revealed a unique activation mechanism characterized by an extensive hydrogen bond network extending from the top of TM7 through TM1, TM2, and down to the ribose binding site. This network, not observed in other AR subtypes, appears to play a role in facilitating the conformational changes associated with A_3_AR activation. With regards to receptor activation, multiple mutations impaired receptor signalling, while other mutations showed ligand-dependent effects on efficacy, particularly with the agonist NECA. Further exploration of the unique A_3_AR activation mechanism, possibly through additional mutagenesis studies, signalling assays, and molecular dynamics simulations is warranted (*94–97*). In addition, structural studies of the A_3_AR bound to a wider range of ligands, including partial agonists and allosteric modulators, will further elucidate the structural basis of ligand efficacy and allosteric modulation (*30, 79*). Together, these insights could be valuable for the design of novel ligands with tailored efficacy profiles (*98, 99*).

Our structures of A_3_AR in complex with different G protein constructs revealed both conserved and divergent features of the receptor-G protein interface. The core interactions involving the last five residues of the G protein α5-helix appear to be consistent across different ligands and G protein constructs. However, the observed differences in the orientation of the α5-helix and αN-helix between the adenosine- and Piclidenoson-bound structures raise intriguing questions about ligand-dependent modulation of G protein coupling. The reduced number of contacts between A_3_AR and G protein in the adenosine-bound structure compared to the Piclidenoson-bound structure suggests a potential link between the extent of ligand-receptor interactions in the orthosteric site and the stability of the receptor-G protein complex (*79*). This observation aligns with the concept of ligand-dependent signaling bias and the observed differences in G protein coupling between adenosine and Piclidenoson suggest the potential for developing ligands with biased signaling properties (*100, 101*). This could be particularly valuable in therapeutic contexts where selective activation of specific signaling pathways is desired (*102–104*). The use of different G protein constructs (DNG_i_, mG_si_), stabilising methods (scFv16, Nb35), and tethering strategies (C-terminal Gα fusion, LgBit-HiBit tethering between the C-terminus and Gα protein) is common in GPCR cryo-EM studies (*65, 67, 81, 105–107*), but necessitates caution in interpretation and is a general limitation of these studies. Investigation of ligand-dependent G protein coupling using consistent G protein constructs and methodologies to clarify the observed differences between adenosine- and Piclidenoson-bound structures will be important for future studies.

This comprehensive study of A_3_AR structure and pharmacology provides a solid foundation for future research and highlights areas that warrant further research. A key challenge in A_3_AR drug development was significant variation in ligand pharmacology across different species of receptor (*74*). The high-resolution structures presented here offer a structural framework for determining the molecular basis of species-dependent differences in A_3_AR pharmacology, which would aid future translation studies. In addition, these structures provide an opportunity to reinterpret existing medicinal chemistry and structure-activity relationship data for A_3_AR ligands (*85, 108–110*). Integration of these structural insights with previous findings will allow for more rational and targeted approaches to ligand design. These advances open new avenues for developing highly selective and efficacious A_3_AR ligands with potential therapeutic applications across a spectrum of disorders, including inflammatory and liver diseases, and glaucoma.

## Materials and Methods

### Materials

FreeStyle 293-F suspension cells, FreeStyle 293 expression serum-free media, Dulbecco’s modified Eagle’s medium (DMEM), fetal bovine serum (FBS), and trypsin were purchased from Invitrogen. Rolipram and forskolin were purchased from Sigma. Guanosine 5’-(γ-thio) triphosphate, [35S]-, 1250Ci/mmol was purchased from PerkinElmer Life. N(6)-(3-iodobenzyl)-5’-N-methylcarboxamidoadenosine (IB-MECA/Piclidenoson), 2-Cl-IB-MECA was purchased MedChem Express (Monmouth Junction, USA). NECA (adenosine-5’-N-ethyluronamide), MRS1220 (9-Chloro-2-(2-furanyl)-5-((phenylacetyl)amino)-[1,2,4]triazolo[1,5-c]quinazoline) and adenosine were purchased from Sigma (St. Louis, MO, USA). Adenosine deaminase (ADA) was purchased from Roche Applied Sciences (Mannheim, Germany). Furimazine was purchased from Promega (Alexandria, Australia) and coelentrazine-h from Nanolight Technology (Arizona, USA). Tested compounds were dissolved in dimethyl sulfoxide. Fluorescent conjugates of xanthine amine congener (XAC) CA200645 were purchased from CellAura Technologies Ltd. (Nottingham, UK). For western blots, mouse poly-His antibody was purchased from QIAGEN, and IRDye 680RD goat anti-Mouse IgG was from LI-COR. Lauryl maltose neopentyl glycol (LMNG) and cholesteryl hemisuccinate (CHS) were purchased from Anatrace. Benzonase was purchased from Merk Millipore, and apyrase was from NEB.

### Cell lines

FreeStyle 293-F suspension cells for transient transfection of WT-A_3_AR, the A_3_AR expression, and A_3_AR 3C construct were grown in FreeStyle 293 expression serum-free media (Gibco) and maintained in a 37 °C incubator containing a humidified atmosphere with 5% CO2 on a shaker platform rotating at 120 rpm. Human HEK 93 adherent (HEK293a) cells for transient transfection of Nluc-A_3_AR were grown in DMEM with 10% (v/v) FBS and maintained at 37 ℃ with 5% CO2. Chinese hamster ovary (CHO) cells stably expressing the human adenosine A3 receptor (Flp-In-CHO cells) were generated as previously described (*111*). Cells were grown in the culture DMEM containing 10% (v/v) FBS and selection antibiotic hygromycin B (0.5 mg/mL) at 37 ℃ with 5% CO2. Insect cells *Spodoptera frugiperda* (Sf9) and *Trichoplusia ni* (TNI) were grown in ESF 921 serum-free media (Expression System) and shaken at 27 °C.

### Constructs

The A_3_AR-BRIL-S97R construct was designed in a pFastBac vector as discussed in the results section. The anti-BRIL Fab (BAG2) and anti-BAG2 nanobody (Nb) were gifts from Christopher Garcia’s laboratory (*43, 112*). The anti-BAG2 Nb was cloned in pET26b+ vector with an N-terminal histidine tag followed by a TEV protease site. For the A3AR G protein-bound complexes, the human wild-type A_3_AR (Uniprot ID: P0DMS8) was modified with a FLAG epitope at the N-terminus and a Histidine tag at the C-terminus to facilitate purification and detection of A_3_AR was cloned into a pFastBac vector. To improve receptor expression, the first 22 amino acids of the human M_4_ muscarinic receptor (M_4_ mAChR), which contains 3 N-glycosylation sites (*32*), were inserted between the FLAG tag and the native A_3_AR sequence. To validate the pharmacological behaviour of the A_3_AR expression construct, we designed constructs that include the above modifications in a pcDNA5 vector for expression in mammalian cells. The adenosine-bound G protein complex was formed using a dominant negative form of Gα_i1_(DNGα_i1_), a dual expression vector containing Gγ_2_ and 8xHis tagged Gβ_1_ (*34, 66, 67*). Four Gα_i1_ subunit mutations in DNG_i1_ alter nucleotide binding and affinity for Gβγ to prevent complex dissociation. The Piclidenoson-bound G protein complex was formed using mini-G_si_ (*68*), which contains a shortened Gα_s_ subunit lacking the α-helical domain (AHD), in which mGα_s_ residues were replaced with Gα_i_ residues close to the receptor-G protein. A GGGS linker was introduced between the carboxyl terminus of A_3_AR and G protein (*107, 113*). Additionally, in the mG_si_ sequence, the αN of mGα_s_ was replaced by those in Gα_i_ (G2HN.02 to K35S1.03). Finally, a 3C cleavage site was incorporated next to the linker prior to the mG_si_ to enable cleavage of the G protein after complex formation. An 8xHis tagged Nb35 in pET20b was provided by B. Kobilka (*69*).

### Cell membrane preparation

WT-A_3_AR and expression A_3_AR constructs were transfected using 1.3 μg DNA per 1 mL of FreeStyle 293F suspension cells at a ratio of 4:1 PEI/DNA and then incubated for 24 hours in a humidified incubator at 37 °C. Transfected cells were harvested and washed with PBS, followed by resuspension in a homogenisation buffer containing 20 mM HEPES, 6 mM MgCl2, 100 mM NaCl, 1 mM EGTA, 1 mM EDTA, pH 7.4. Cell pellets were again resuspended in the homogenisation buffer after centrifugation at 300 g for 3 mins at 4 °C. Resuspended cell mixtures were applied to a homogeniser with three bursts for 10 seconds on ice. Cell nuclei were removed by centrifugation at 600 g for 10 mins at 4 °C. The resulting supernatants were further centrifuged at high speed (30000 g, 60 mins, 4 °C). Cell membranes were collected from pellets and resuspended in a homogenisation buffer. The protein concentrations were determined using bicinchoninic acid (BCA) protein assay. Cell membranes were stored at −80 °C.

### [^35^S]-GTPγS binding assay

Adenosine-5’-N-ethyluronamide (NECA)-mediated binding of [^35^S]-GTPγS was used to measure G-protein activation by the receptors in membranes extracted from transfected FreeStyle HEK293F cells. Experiments were performed as described previously (*34*).

### Nano-BRET binding assay

HEK293a cells were seeded into 10 cm dishes at a density of 4 million cells per dish. Following an eight-hour incubation, cells were transfected with 5 μg of N-terminal Nluc-tagged A_3_AR constructs, using polyethylenimine (PEI) at a 4:1 PEI-to-DNA ratio. After 24 hours, the cells were replated into 96-well, white-bottom, poly-D-lysine-coated culture plates at a density of 40,000 cells per well. Nano-BRET saturation binding was performed as previously reported (*51, 52*) using the fluorescent A_3_AR antagonist CA200645 (final concentration ranging from 15 nM to 1000 nM). Before the assay, cell media was replaced with BRET buffer (HBSS supplemented with 10 mM HEPES, 1 unit/mL of adenosine deaminase (ADA), and 10 µM MRS1220 was used to define nonspecific binding. Following equilibration at 37 ℃ in a humidified atmosphere for 50 mins, the substrate 10 µM furimazine or coelenterazine-h was added to each well and cells incubated for a further 10 minutes at 37 ℃ in a humidified atmosphere. Bioluminescence emission wavelengths were measured as previously described (*114*) using a PheraSTAR Omega plate reader (BMG Labtech) using 460 nm (80 nm bandpass; donor NanoLuc emission) and >610 nm (long pass filter; fluorescent ligand emission). The raw BRET ratio was calculated by dividing the >610 nm emission by the 460 nm emission. All experiments were performed in duplicate. Competition binding experiments were performed similarly using a concentration of CA200645 near the K_d_ (200 nM) of the A_3_AR construct and 10-fold serial dilutions of competitors with an equilibrium time of 50 minutes. To determine the irreversible binding between LUF7602 and Nluc-tagged hA_3_AR-BRIL-S97R, we assessed the competitive binding capacity of LUF7602, PSB11, and MRS1220 to two groups of cells (washed and unwashed) at a concentration of 10 μM with 4 hours of incubation at room temperature. The washed group was subjected to 4 washing steps with BRET buffer (10 mins intervals) to remove unbound ligands. Subsequently, 100 nM of CA200645 was added to both groups and incubated for 50 minutes before adding furimazine. In the site-directed mutagenesis study, 200 nM of CA200645 was used.

### Trupath assay

HEK293a cells were seeded into 96-well, white-bottom, poly-D-lysine-coated culture plates at a density of 20,000 cells per well. After four hours of incubation, cells were transiently transfected with equal parts of N-terminal Nluc-tagged A_3_AR constructs and G proteins (Gi1-Rluc8, Gβ3, and Gγ9-GFP2), using 15 ng of each DNA construct for a total of 60 ng of DNA per well. Polyethylenimine (PEI) was added at a 6:1 PEI-to-DNA ratio. Two days post-transfection, plates were washed twice with BRET buffer (HBSS with 10 mM HEPES, pH 7.4). After a 30-minute incubation at 37 ℃, plates were placed in a PHERAstar plate reader at 37℃, and four baseline readings were taken using BRET2 filters at 410 ± 80 nm and 515 ± 30 nm. Increasing concentrations of adenosine, NECA, and CF101 were then added, followed by eight additional readings. BRET2 ratios were calculated as the ratio of GFP2 emission (515 ± 30 nm) to RLuc8 emission (410 ± 80 nm).

### Data and Statistical Analysis

Data were expressed as the mean ± standard error. All binding and Trupath data were analysed using the nonlinear regression curve-fitting program with Prism 10 (GraphPad, San Diego, CA, USA). For nano-BRET saturation binding experiments, specific binding was calculated by subtracting non-specific binding (defined as binding in the presence of 10 µM MRS1220) from total binding. Specific binding data in each experiment were then normalised to the maximal binding (B_max_) of WT and fit to a one-site specific binding equation: Y= B_max_*[X]/(K_d_+[X]). For nano-BRET competition binding experiments, data were normalised total and non-specific binding responses and fit to a one site – Fit K_i_ equation. Trupath experimental data was baseline-corrected to the initial four baseline reads and then adjusted to the buffer control response. The area under the curve (AUC) for each response was calculated and plotted with a non-linear regression curve (log[agonist] vs. response, three-parameter model), using the equation: Y=Bottom + (Top-Bottom)/(1+10^((LogEC50-X))). Here, X represents the log dose of the agonist, Y is the BRET2 AUC response, Top and Bottom are the curve plateaus, and EC_50_ is the agonist concentration yielding a response halfway between the Bottom and Top values. To determine efficacy values. The efficacy of the agonists (1_A_) were separately calculated using the Black–Leff operational model of agonism (*77*) using K_A_ values from **Table 2** and then corrected for receptor expression relative to WT using B_max_ values from **Table 1** (*78*).Significant differences versus WT were determined by one-way ANOVA with a Dunnett’s multiple comparison test. Significance levels are indicated as follows: p ≤ 0.0001 (****), p = 0.0001–0.001 (***), p = 0.001–0.01 (**), and p = 0.01–0.05 (*).

### Receptor and G protein expression

The Bac-to-Bac Baculovirus Expression System (Invitrogen) produced high-titre recombinant baculovirus for A_3_AR constructs and DNGα_i1_. BestBac linearized DNA (Expression Systems) was used to make baculovirus for Gβ_1_γ_2_. To make the adenosine-bound A_3_AR-G_i_ complex, *Trichoplusia ni* (Tni) cells were co-infected with A_3_AR, DNGα_i1_, and Gβ_1_γ_2_ baculovirus at the multiplicity of infection (MOI) ratio of 4:2:1 at a density of 3.5 ∼ 4 million/mL. To make the Piclidenoson-bound A_3_AR-mG_si_ complex, a 2:1 ratio of A_3_AR-mG_si_:Gβ_1_γ_2_ baculovirus was used. The A_3_AR-BRIL-S97R construct was expressed in *Spodoptera frugiperda* (Sf9) cells. Insect cell cultures were shaken at 27 ℃ in ESF 921 serum-free media.

### Expression and purification of scFv16 and Nb35

ScFv16 was expressed and purified as previously described (*65*). The expression and purification of Nb35 were adapted from prior methods (*69*). In brief, Nb35 was expressed in BL21(DE3) Rosetta *E. Coli* strain using the autoinduction method followed by purification as previously described.

### Purification of A_3_AR-G protein complexes

The purification of A_3_AR-G protein complexes was performed as previously described (*34*) with minor modifications. The High 5 cell pellet was thawed and lysed in a hypotonic buffer containing 20 mM HEPES pH 7.4, 50 mM NaCl, 2 mM MgCl2, 10 μM agonist, protease inhibitors (200 μM phenylmethylsulfonyl fluoride (PMSF), 1 mM leupeptin and trypsin inhibitors (LT), 0.2 mg/ml benzamidine), 1 mg/ml iodoacetamide, 50 μg/mL, 2.5 units of apyrase (NEB) and benzonase (Merk Millipore) and stirred at 25 ℃ for 20 mins to obtain homogeneity. The cell lysate was centrifuged at 20,000 g for 20 mins at 4 ℃. Cell membranes were resuspended and homogenised with a dounce homogenizer in buffer containing 30 mM HEPES pH 7.4, 100 mM NaCl, benzonase (Merk Millipore, 2 ul/800 ml), 2.5 units of apyrase (NEB), protease inhibitors (200 μM PMSF, 1 mM LT, 0.195 mg/ml benzamidine), 2 mM CaCl2, 2 mM MgCl2, and 10 μM agonist in the presence of 0.5 % (w/v) lauryl maltose-neopentyl glycol (LMNG) buffer 0.03% (w/v) cholesterol hemisuccinate (Anatrace, CHS) for 2 hours at 4 ℃. After removal of insoluble debris by centrifugation (30,000 g for 30 minutes, 4 ℃), the supernatant was then loaded onto a glass column filled with M1 anti-Flag antibody resin and washed with 20 column volumes (CV) of wash buffer containing 30 mM HEPES pH 7.5, 100 mM NaCl, 2 mM CaCl2, 2 mM MgCl2, 10 μM agonist, 0.01 (w/v) % LMNG, and 0.001 % (w/v) CHS. Receptor complex was eluted from the anti-FLAG resin using the wash buffer supplemented with 0.2 mg/ml flag peptide and 10 mM EGTA. Excessive scFv16 (for DNGαi1) or Nb35 (for mGsi) was added to the eluate at a 1:1.5 molar ratio for 30 mins at 4℃. The receptor complex was then concentrated using an Amicon Ultra Centrifugal Filter Unit (MWCO, 100 kDa), filtered (0.22 *μ*m), and subjected to size-exclusion chromatography (SEC) using a Superdex S200 increase 10/300 column (GE Healthcare) in buffer containing 30 mM HEPES pH 7.5, 100 mM NaCl, 10 μM agonist, 0.01 % LMNG and 0.001 % CHS. Monodisperse fractions of all the components were pooled together and spiked with 10 μM agonist before concentrating and being flash-frozen in small aliquots in liquid nitrogen and stored at −80°C. Protein complexes were confirmed by Coomassie-stained SDS-PAGE gels, detection of FLAG and His epitopes by Western Blot, and by negative stain EM.

### Expression and purification of BAG2, elbow Nb, A_3_AR-BRIL-S97R, and complex assembly

The BAG2, anti-BRIL Fab fragment, and elbow Nb were expressed in *E. Coli* BL21 (Gold) and purified as previously described (*43*). Purification of A_3_AR-BRIL-S97R was performed in the absence of ligands with adenosine deaminase added to remove endogenous adenosine. The purification of A_3_AR-BRIL-S97R was similar to that of previous methods. Expressed cell pellets were lysed by osmotic shock followed by solubilization in detergent buffer (1% DDM and 0.03% CHS). Solubilised receptor was batch-bound to Ni-chelating resin, followed by washing, and elution. The eluted sample was mixed with anti-Flag M1 antibody resin and loaded onto a glass column. The receptor was buffer exchanged into the detergent 0.1% LMNG with 0.01% CHS and eluted. The receptor was concentrated and purified further by SEC. Purified A_3_AR-BRIL-S97R was then incubated with 30 μM LUF7602 and excess BAG2 and elbow Nb at a 1:2:4 molar ratio for 2 hours at room temperature and overnight at 4 °C. The LUF7602-bound A_3_AR-BRIL-S97R-BAG2-Nb complex was purified by SEC. Fractions containing the full complex were collected and spiked with LUF7602 at a final concentration of 10 µM and incubated on ice for 1 hour before concentrating to 12 mg/mL. The protein complex was confirmed by Coomassie-stained SDS-PAGE gels, detection of FLAG and His tags by Western Blot, and by negative stain EM.

### Cryo-EM sample preparation and data collection

Adenosine-bound A_3_AR-DNG_i_-scFv16 cryo-EM grids were prepared on UltrAuFoil 300 mesh 1.2/1.3 grids (Quantifoil, Au300-R1.2/1.3). A_3_AR-mG_si_-Nb35-IB-MECA and A_3_BRIL-S97R-BAG2-Nb-LUF7602 cryo-EM grids were prepared on EMAsian - TiNi 200 mesh 1.2/1.3 grids. Grids were glow discharged for 180 seconds (UltrAuFoil grids) or 90 seconds (EMAsian) at 15 mA current using Pelco EasyGlow in low-pressure air. 3 μL of the purified samples were applied to each grid. Excess sample on the grids was removed by blotting on an FEI Vitrobot Mark IV (Thermo Fisher Scientific) at 100 % humidity and 4 °C with a Whatman #1 filter paper, with a blot force of 12 and blot time of 2s for UltrAuFoil grids and a blot force of 4 for blot time of 2s for the EMAsian grids. Blotted grids were then vitrified by plunging into liquid ethane and then liquid nitrogen.

For the adenosine-bound A_3_AR sample, cryo-EM imaging was performed on the Arctica microscope (Thermo Fisher Scientific) operating at 200 kV equipped with a Gatan K2 Summit direct electron detector in counting mode, corresponding to a pixel size of 1.03 Å. A total of 8676 movies were collected using beam shift with the 9-hole acquisition. Cryo-EM imaging of A_3_AR-mGs_i_-Nb35-IB-MECA was performed on a Titan Krios G3i transmission electron microscope (Thermo Fisher Scientific) operated at an accelerating voltage of 300 kV at a nominal magnification of 105000 in nanoprobe EFTEM mode. Gatan K3 direct electron detector and a 50 μm C2 aperture without an objective aperture inserted during the data collection period were applied to acquire dose-fractionated images of the samples with a slit width of 10 eV, pixel size 0.82 Å. For each movie stack, 60 frames were recorded with a total exposure dose of 60 e-/A2. In total, 6526 movies were recorded in super-resolution mode using SerialEM. For A_3_BRIL-S97R-BAG2-Nb-LUF7602, 6398 movies were collected on the Titan at a magnification of 130000x in nanoprobe TEM mode, with electron counting mode with a physical pixel size of 0.65 Å/pixel, exposure rate of 10.57 counts per pixel per second, exposure time of 2.68 s and a total dose of 60 e /Å2.

### Image processing

Data processing for the A_3_AR-DNG_i1_-scFv16-adenosine complex was performed with the movie stacks subjected to beam-induced motion correction using UCSF MotionCor2 (*115*) using 5 × 5 patches. Contrast transfer function (CTF) parameters for each corrected micrograph were calculated by Gctf (*116*). Images with Gctf maximum resolution estimates worse than 4 Å were excluded from further processing. Automated particle selection yielded 7274091 particles using the Gautomatch software package. The particles were imported into Relion 3.1 for binned extraction with a box size of 240 pixels and re-scaled to 60 pixels. The extracted particles were carried to cryoSPARC (v3.1) software package (*117*) and subjected to 2D classification and *ab initio* 3D and 3D refinement to select good particles, yielding ∼1.2 M particles. Particles were taken to Relion3.1 for Bayesian particle polishing, CTF refinement, 3D auto refinement and subsequent 3D refinement back in Cryosparc, producing a final map at 2.9 Å resolution from 325,569 particles. The resulting particle set was then subjected to 3D refinement and post-processed with a mask excluding the detergent micelle and G proteins, resulting in the final receptor-focused map at a resolution of 3.44 Å according to the FSC 0.143 criteria. The local resolution was estimated using the cryoSPARC v2.15 local resolution estimation function.

For the A_3_AR-mG_si_-Nb35-IB-MECA complex, all of the processing was performed in cryoSPARC. A total of 6526 micrographs were motion corrected using Patch motion correction, and CTF parameters were estimated using patch CTF estimation. Images with the highest resolution of less than 4 Å were selected for further processing. A total of 5876 movies were finally chosen for particle picking. Particles were first picked using the template picker, followed by 2D classification. Good 2D class averages with randomised orientations produced a particle stack of 2407599 particles (from a total of 6.0M extracted particles). The good particles and 203651 bad-looking particles were used to generate two *ab initio* 3D models. The initial models were then used as the templates for initial 3D refinement, followed by another round of 2D classification on particle set matching features of the GPCR heterotrimeric complex. The subsequent selected 2D averages, containing 943812 particles, were applied to another round of 2D classification and hetero refinement using the same initial models, resulting in a final particle set for further processing. Finally, 589921 particles were selected for a non-uniform 3D consensus refinement and CTF envelope fitting, yielding a map with a resolution of 2.6 Å at a Fourier shell correlation of 0.143 (gold standard). The dataset was subjected to further local-motion correction (per particle), and the final resolution was improved to 2.5 Å. Subsequently, to further improve resolution in the transmembrane region, a mask excluding the detergent micelle and G proteins was implemented to calculate a high-quality map of the receptor, producing a receptor-focused density map of resolution at 3.1 Å, which was used to build the receptor model.

For the A_3_BRIL-S97R-BAG2-Nb-LUF7602 complex, movies were motion-corrected using Patch Motion Correction in CryoSPARC (v3.1). Patch CTF was used to estimate defocus values. Micrographs with CTF resolution estimates worse than 4 Å were excluded from further examination. The ∼3.3M particles were auto-picked using a previously generated template. The particles were extracted with a box size of 360 pixels re-scaled to 90 pixels. The extracted particles were subjected to 2D classification, *ab initio* 3D, and heterogeneous 3D refinement. Particles were re-extracted at full resolution and imported into Relion 3.1 for Bayesian particle polishing and CTF refinement. The shiny particles were imported into the CryoSPARC (v3.1) for additional 2D and 3D classifications. Final set of 328,104 yielded a final map at 2.6 Å. For A_3_AR, a local class for the receptor-only region was performed with a mask excluding the BRIL-BAG2-Nb to improve the map quality for the ligand binding pocket to assist modelling, resulting in the final consensus map at 3.3 Å according to the FSC 0.143 criteria.

### Model building and refinement

#### A_3_AR-DNG_i1_-scFv16-adenosine complex modelling

An initial homology model of A_3_AR was generated using the SWISS-MODEL server (*118*) with the activated structure of A_1_AR (PDB: 7LD4) as a template (*36*). Models of G_i_ heterotrimers and scFv16 were adopted from dopamine receptor D_3_R-Gi-Pramipexole complex (PDB: 7CMU) (*119*). A resembled model was docked into the cryo-EM map using the “fit in map” routine in UCSF Chimera (*120*). This starting model was then subjected to rigid body fitting and followed by iterative rounds of automated refinement in Phenix real-space refinement (*121*) and repeated rounds of manual building in Coot (*122*).

#### A_3_AR-mG_si_-Nb35-IB-MECA complex modelling

The receptor from the model of A_3_AR-DNG_i1_-scFv16-adenosine was rigid-body placed into the receptor-focused cryo-EM map. The mG_si_, Gβ_1_γ_2_, and Nb35 were modified from the adhesion GPCR ADGRD1 structure (PDB: 7WU2) (*123*) and were rigid-body placed in the consensus map. The fitted models of all subunits were further refined for several iterations of manual model building in COOT and real-space refinement, as implemented within the Phenix software, respectively.

#### A_3_AR-BRIL-S97R-BAG2-Nb-LUF7602 complex

A homology model of A_3_AR generated by SWISS-MODEL using the A_1_AR crystal structure (5UEN) (*32*) as a template was placed into the receptor-focused map in ChimeraX, followed by repeated rounds of model building in Coot and iterative refinement with Phenix. The BRIL-BAG2-Nb trimer module was obtained from the cryo-EM structure of the Frizzled complex (PDB:6WW2) (*42*) and was docked into the consensus cryo-EM density map using ChimeraX and rigid-body placed with Phenix. Refinements with Phenix and manual building in Coot were performed on the trimer. To aid with molecular modelling composite maps were made from DeepEMhancer post-processing cryo-EM maps of the consensus and receptor focused refinement maps (*124*). The molecular model was refined against the composite map (EMD-48065). All the final models were visually inspected for the general fit to the map and validated using Molprobity (*125*). Structural figures were prepared using ChimeraX (*126*). The cryo-EM data collection, refinement and validation report for this complex are shown in **Table S1**.

### Induced fit docking (IFD)

The induced fit docking (IFD) protocol of Schrödinger was used to dock XAC into A3. This protocol uses a combination of Glide XP and Prime to accurately model ligand and protein side chains. The protein was prepared using the Schrödinger Protein Preparation Wizard. This process adds hydrogen atoms, assigns bond orders, optimises the hydrogen-bonding network, and performs a restrained minimisation using the OPLS4 force field. XAC was prepared from SMILES using Schrödinger LigPrep, to generate 3D structures, different ionisation states and tautomeric forms at a physiological pH of 7.4. The ligand structures were minimised using the OPLS4 force field. The grid box was built with the centroid of the C59 ligand, Prime refinement of side chains within 5 Å, and a maximum of 20 poses were redocked with Glide XP.

## Acknowledgments

The AI tool Claude 3.5 Sonnet (Anthropic, 2024) was used for proofreading of the manuscript checking for consistencies in spelling, grammar, and clarity of the text. It was not used for the generation of ideas, content, figures, or data.

## Contributions

Conceptualization: LZ, LTM, AC, AG, DMT

Methodology: LZ, JIM, ATN, DE, LHH, LTM, AG, DMT

Investigation: LZ, JIM, FMB, HV, FMB, AG

Funding acquisition: DMT

Supervision Writing – original draft: LZ, JIM, DMT

Writing – review & editing: All authors

## Data and materials availability

All data needed to evaluate the conclusions in the paper are present in the paper and/or the Supplementary Materials.

