## Supplemental Materials for "Molecular basis of ligand binding and receptor activation at the human A3 adenosine receptor"

**Affiliations:**
[1]Drug Discovery Biology, Monash Institute of Pharmaceutical Sciences, Monash University; Parkville, Australia.
[2]ARC Centre for Cryo-electron Microscopy of Membrane Proteins, Monash Institute of Pharmaceutical Sciences, Monash University; Parkville, Australia.
[3]Ramaciotti Centre for Cryo-Electron Microscopy, Monash University; Clayton, Australia.
[4]Neuromedicines Discovery Centre, Monash University; Parkville, Australia.
[5]Division of Medicinal Chemistry, Leiden Academic Centre for Drug Research, Leiden University; Leiden, The Netherlands.
[6]Oncode Institute; Leiden, The Netherlands
[7]The Walter and Eliza Hall Institute of Medical Research; Parkville, Australia
[8]Department of Biochemistry and Pharmacology, The University of Melbourne; Melbourne, Australia
[†]These authors contributed equally to this work.

**The PDF file includes:**

Figs. S1 to S8
Tables S1 to S2

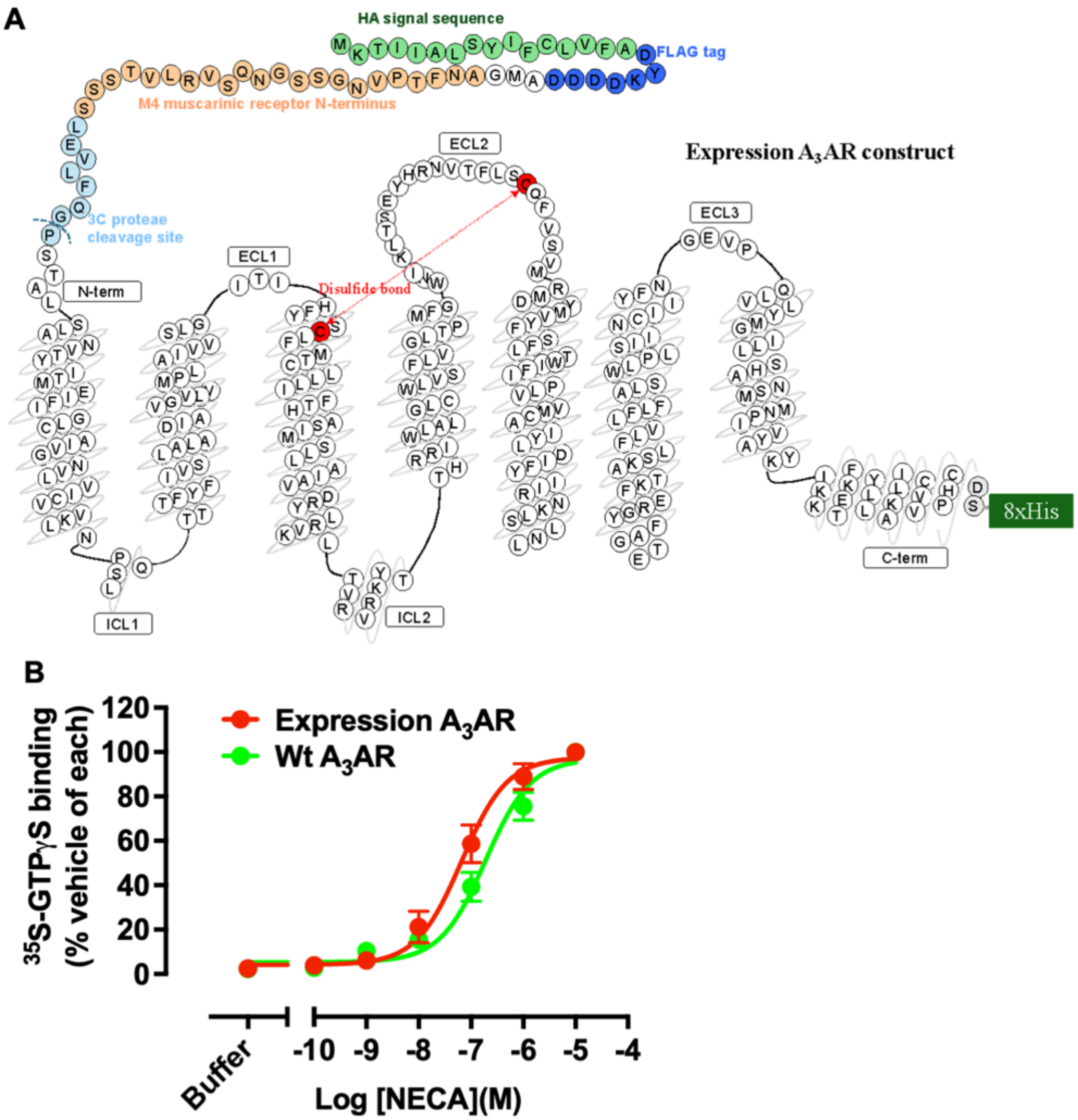


**Fig. S1. Schematic and pharmacology of the A3AR expression construct used in this study.** **(A)** Snake plot of the A3AR expression construct highlighting key features including a N-terminal FLAG epitope followed by the M4 mAChR N-terminus, a 3C protease cleavage site and a C-terminal 8X-His tag.

**(B)** Comparison of [$^{35}$S]-GTPγS binding between the expression A3AR and WT A3AR constructs. Data represent the mean ± SEM from n=3 experiments performed in quadruplicate. The potency values (pEC50) were 6.7 ± 0.1 (n=3) for WT and 7.2 ± 0.1 (n=3) for the expression A3AR construct. The difference was statistically significant (P value = 0.03) using an unpaired t-test; however, receptor expression was not considered in these experiments, which could affect the potency values.

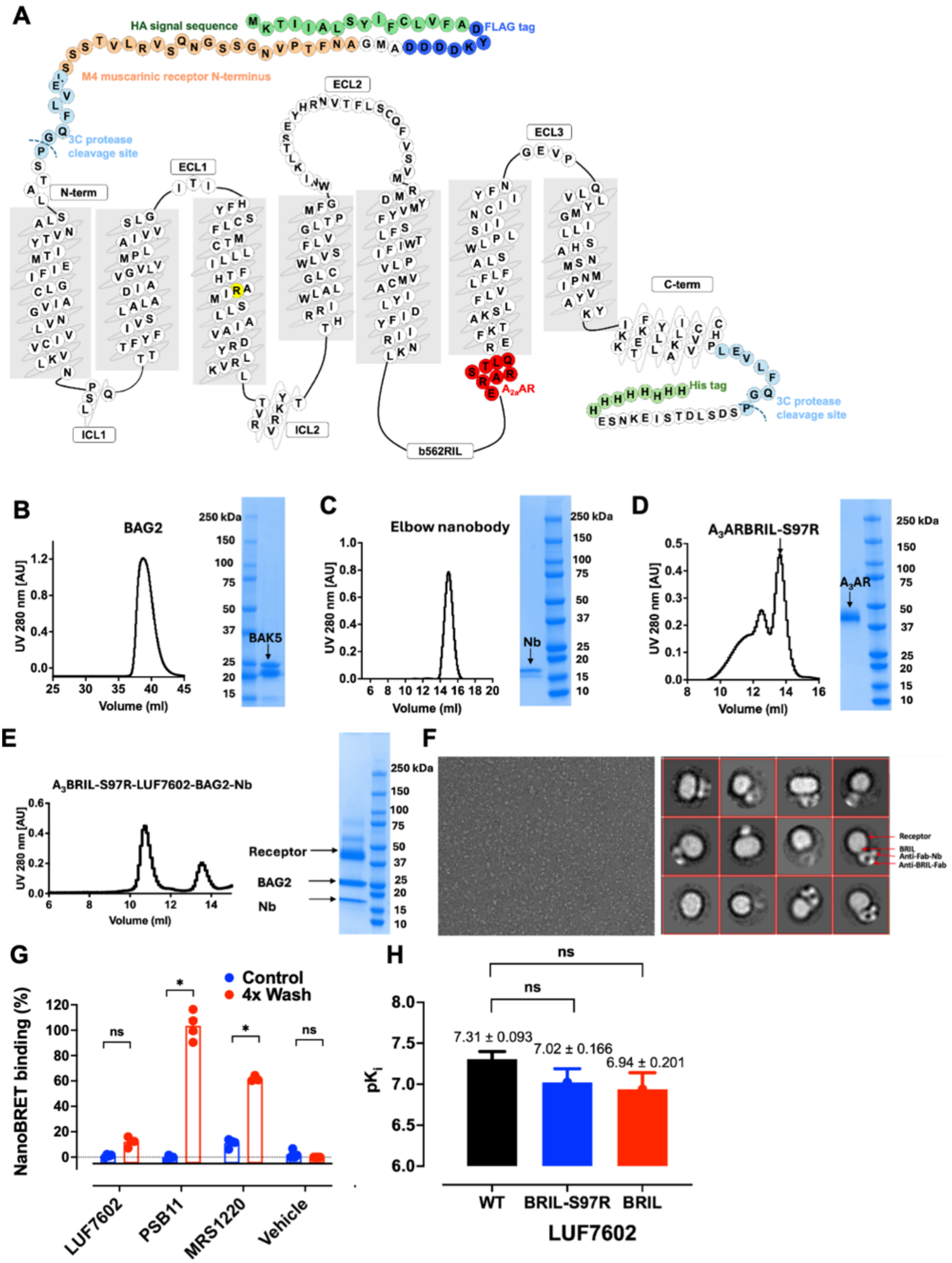


**Fig. S2. Purification of the A3AR-BRIL-S97R-LUF7602-BAG2-Nb complex.**
**(A)** Schematic representation of A3AR construct used to determine the inactive cryo-EM structure, which includes replacing ICL3 with BRIL and 8 residues of TM6 with the equivalent A2AAR residues.

**(B-E)** Size-exclusion chromatography profiles and corresponding SDS-PAGE gels for the purification of (B) the BAG2 chaperone, (C) the elbow nanobody (Nb), (D) $A_3$AR-BRIL-S97R, and (E) the $A_3$BRIL-S97R-LUF7602-BAG2-Nb complex.
**(F)** Negative-stain electron microscopy images of the purified receptor complex showing a raw micrograph on the left and 2D class averages on the right.
**(G)** Inhibition of XAC-630 in a NanoBRET binding assay comparing control (blue) and 4x wash (red) conditions for different antagonists (LUF7602, PSB11, MRS1220, and Vehicle). LUF7602 retained inhibition versus XAC-630 after a 4x washout indicating irreversible binding. Data represent the mean ± SEM from n=3 experiments performed in triplicate. Statistics were performed using paired t-tests: ns = no significant difference; * = $P < 0.05$.
**(H)** Bar graph showing $pK_i$ values for WT, BRIL-S97R, and BRIL constructs with LUF7602. No significant differences (ns) were determined by one-way ANOVA (Prism 10.3.1) with a Dunnett's multiple comparison test.

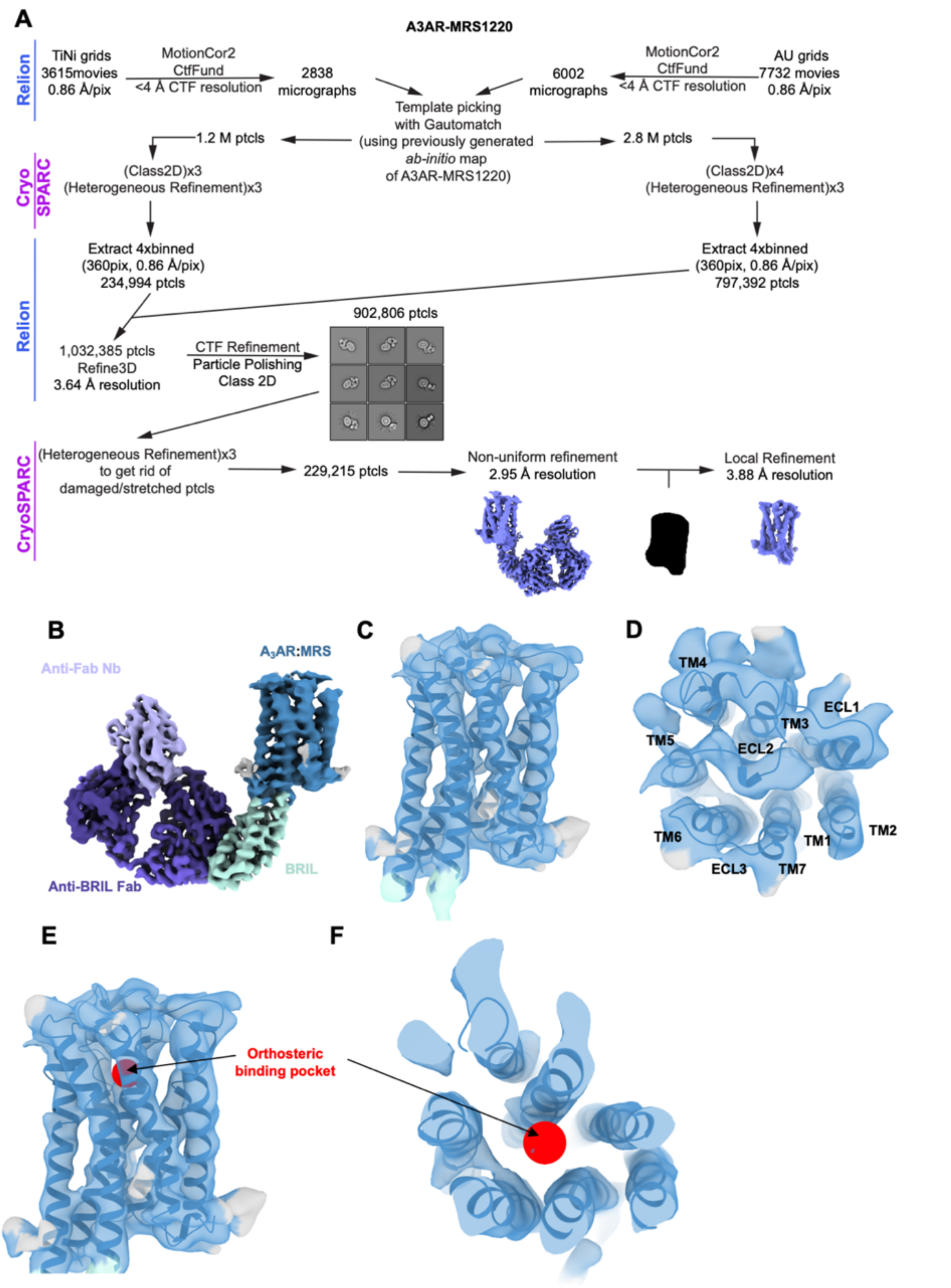


**Fig. S3. Cryo-EM processing and EM map of the $A_3$AR-BRIL-S97R-BAG2-Nb complex purified with MRS1220.**
**(A)** Cryo-EM data processing workflow for MRS1220-$A_3$AR
**(B)** 3D reconstruction of the $A_3$AR-BRIL-S97R-BAG2-Nb, showing the receptor (dark blue), BRIL fusion protein (cyan), anti-Fab nanobody (light purple), and anti-BRIL Fab (dark purple).

**(C)** Cryo-EM density map of the $A_3AR$, showing the overall structure of the receptor and quality of the map.
**(D)** Top view of the $A_3AR$ EM map, highlighting the transmembrane helices (TM1-TM7) and extracellular loops (ECL1-3). ECL2 and the C-terminal portion of ECL1 could not be accurately modelled due to lower resolution surrounding this area.
**(E)** Side view and **(F)** top view of the $A_3AR$ EM map with the adenosine binding site indicated by a red sphere. There was no EM density in the orthosteric binding pocket, which could be due to either MRS1220 not being bound or insufficient resolution to resolve the binding site.

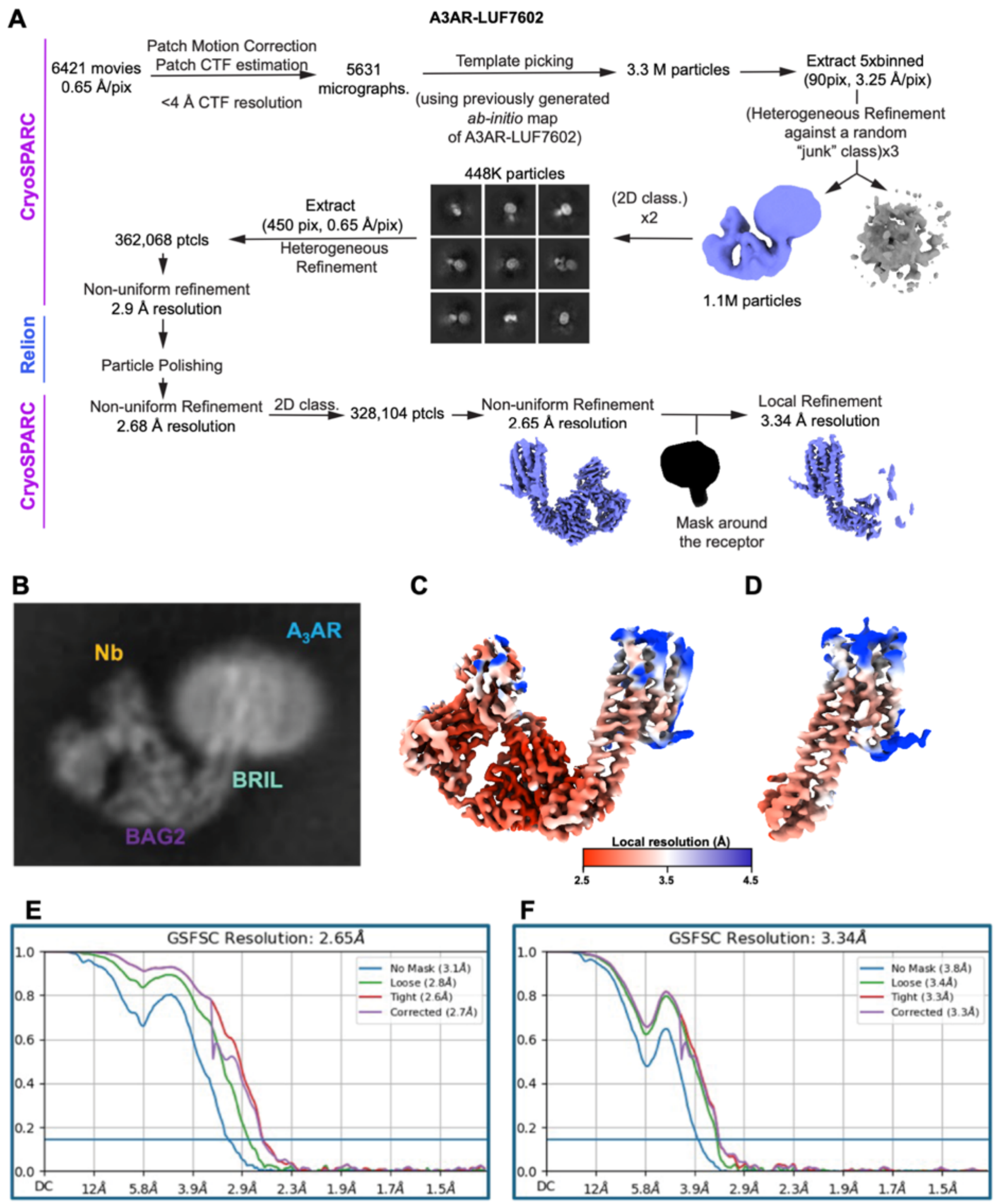


**Fig. S4. Cryo-EM processing and EM maps of the LUF7602-bound to the $A_3$AR-BRIL-S97R-BAG2-Nb complex.**

**(A)** Cryo-EM data processing workflow for LUF7602-$A_3$AR

**(B)** A 2D-class average of the complex in a detergent micelle viewed from the side.

**(C-D)** Local resolution of the (C) consensus map and (D) the local-refined receptor map.

**(E,F)** Gold-standard Fourier shell correlation (GSFSC) plot for (E) the consensus map and the (F) local-refined receptor map.

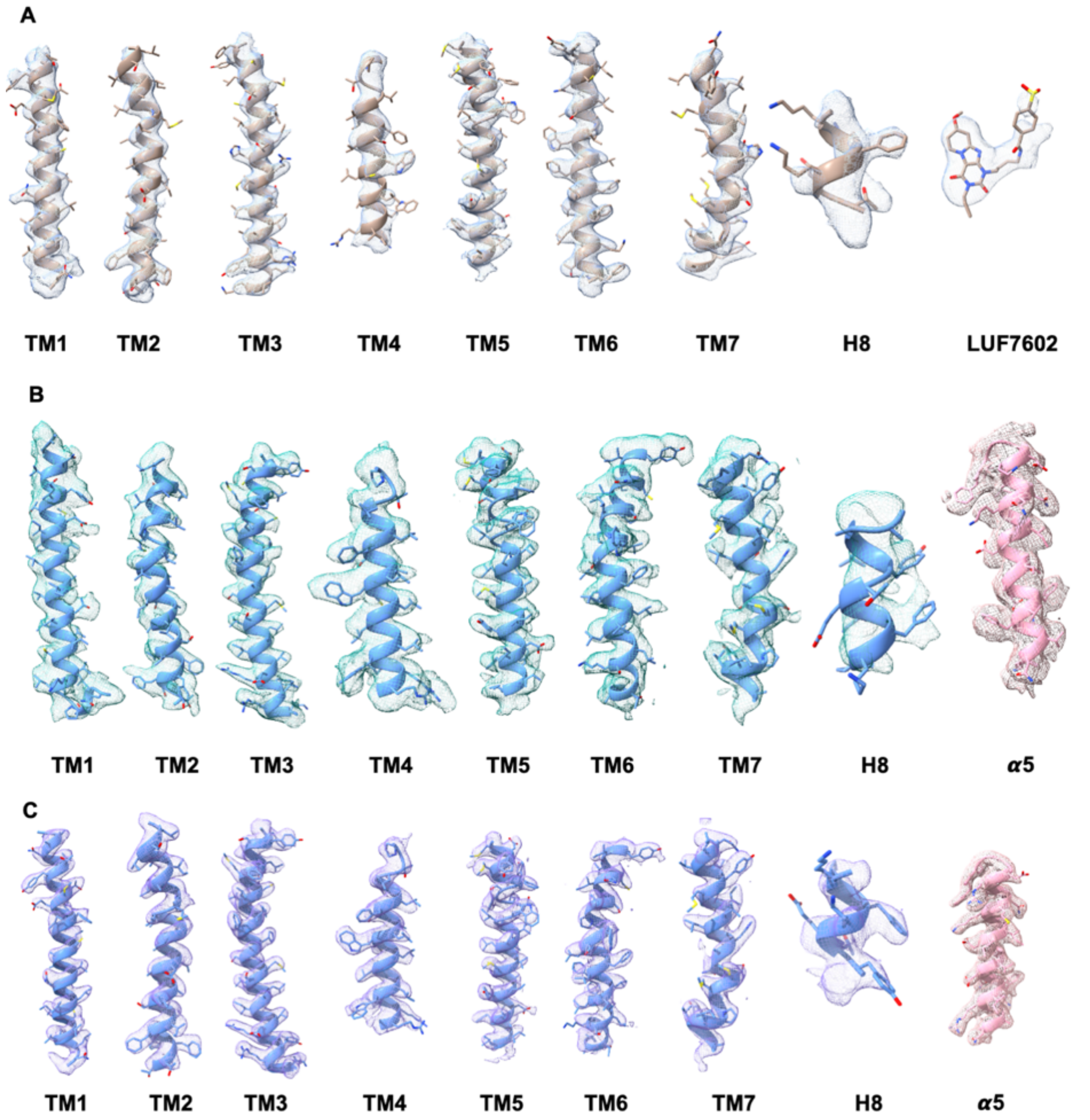


**Fig. S5. Cryo-EM Density Maps**

**(A-C)** Cryo-EM density maps of the seven transmembrane domains (TM1-TM7), helix 8 (H8) for the (A) LUF7602-bound, (B) adenosine-bound, and (C) Piclidenoson-bound $A_3AR$ complex structures. Panels (B,C) include the α5 helix of the G protein.

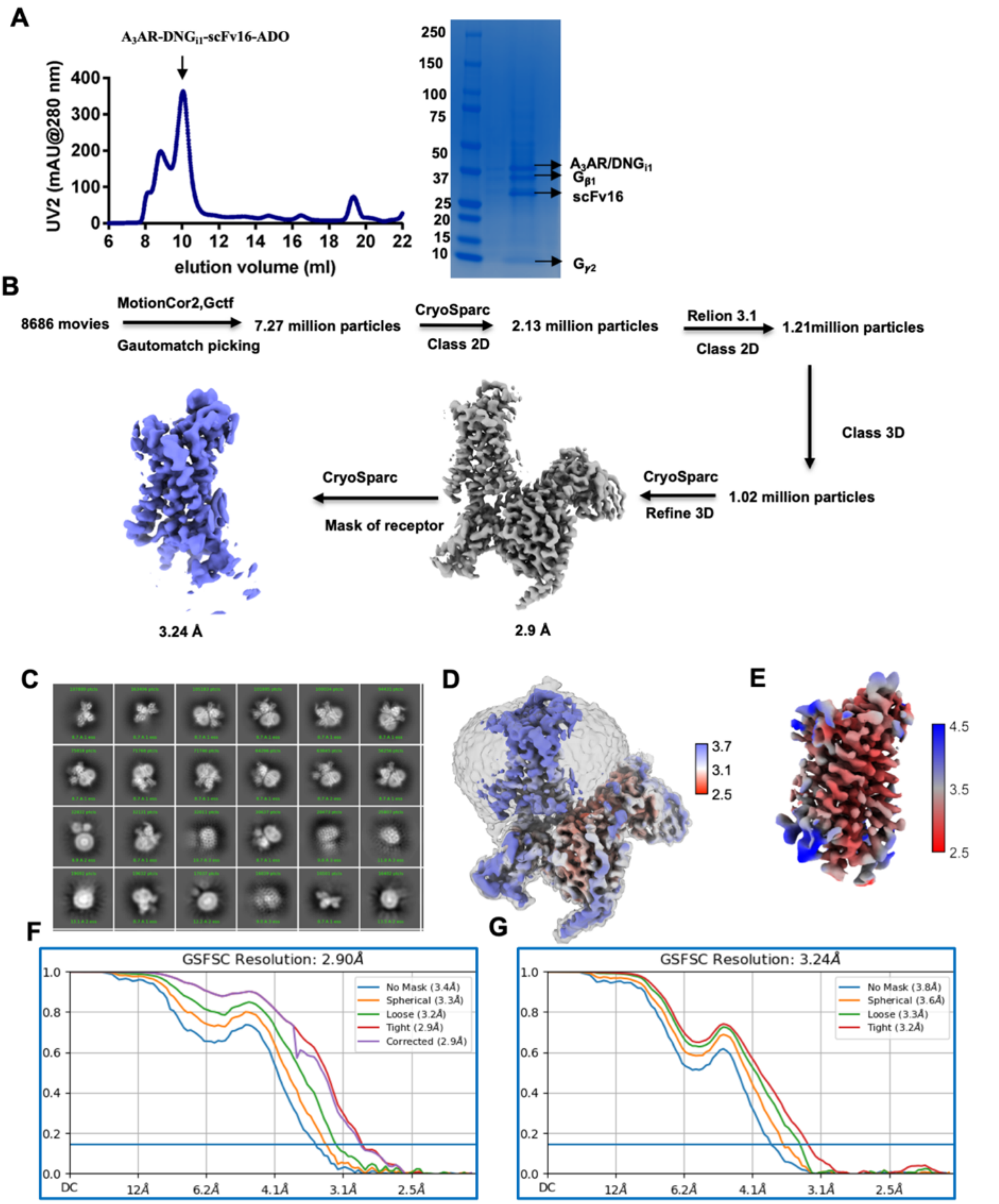


**Fig. S6. Purification and cryo-EM processing of the adenosine-bound $A_3AR$ complex.**
**(A)** Size-exclusion chromatography profile of purified $A_3AR$-$DNG_{i1}$-scFv16-ADO complex and SDS-PAGE gel showing the purified complex components.
**(B)** Cryo-EM data processing workflow for adenosine-$A_3AR$
**(C)** Reference-free 2D class averages of the complex particles.
**(D)** Local resolution of the consensus map ranging from 2.5 Å (red) to 3.7 Å (blue).
**(E)** Local resolution of the local-refined receptor map ranging from 2.5 Å (red) to 4.5 Å (blue).
**(F,G)** Gold-standard Fourier shell correlation (GSFSC) plot for (F) the final consensus map and (G) the local-refined receptor map.

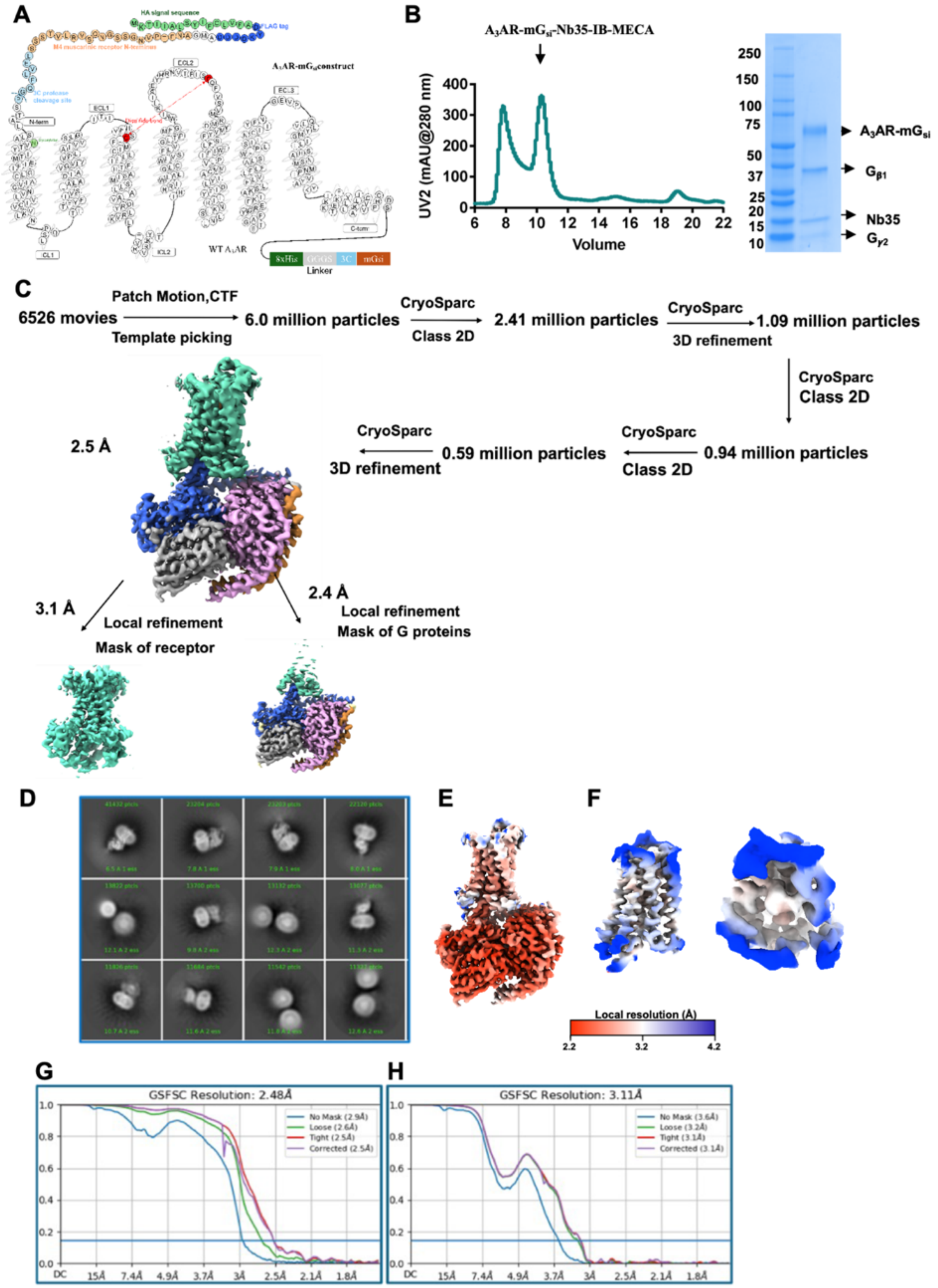


**Fig. S7. Purification and cryo-EM processing of the Piclidenoson-bound A3AR complex.**
**(A)** Schematic representation of the A3AR-mG$_{si}$ construct.
**(B)** Size-exclusion chromatography profile of purified A3AR-mG$_{si}$-Nb35-Piclidenoson complex and SDS-PAGE gel showing the purified complex components.
**(C)** Cryo-EM data processing workflow for Piclidenoson-A3AR.

**(D)** Reference-free 2D class averages of the complex particles.
**(E)** Local resolution of the consensus map ranging from 2.2 Å (red) to 4.2 Å (blue).
**(F)** Two views of the local resolution of the local-refined receptor map ranging from 2.5 Å to 3.5 Å.
**(G,H)** Gold-standard Fourier shell correlation (GSFSC) plot for (G) the final consensus map and (H) the local-refined receptor map.

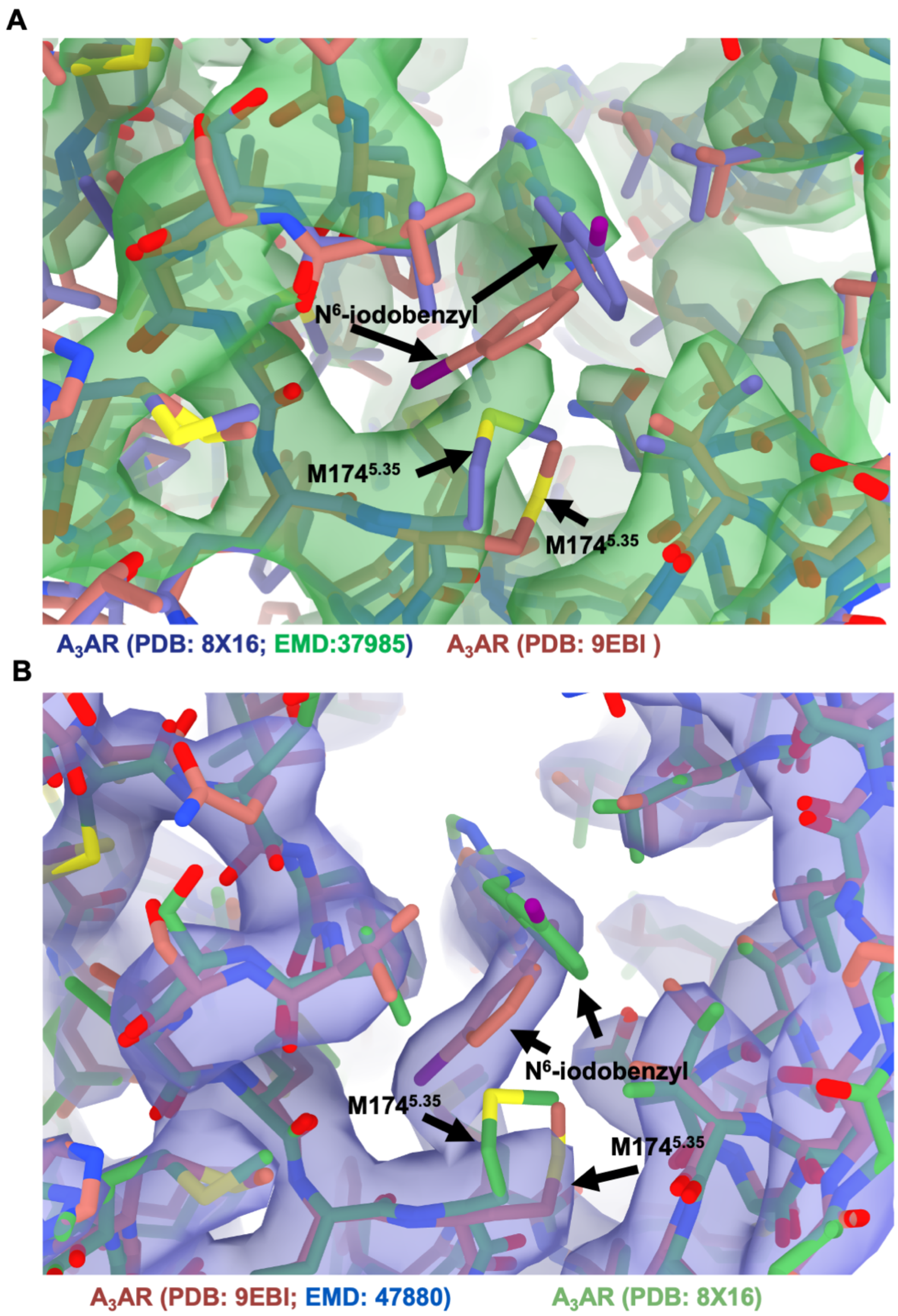


**Fig. S8. Comparison of Piclidenoson-bound A3AR structures**

**(A-B)**. Structural comparison of two $A_3AR$ structures bound to Piclidenoson. The model of PDB: 9EBI is coloured dark red in both panels. (A) The model of PDB: 8X16 is coloured blue and the cryo-EM map is coulored green (EMD:37985). (B) The model of PDB: 8X16 is coloured green and the cryo-EM map for PDB: 9EBI (EMD:47880) is coulored blue. The $N^6$-iodobenzyl group of Piclidenoson and key residue $M174^{5.35}$ is highlighted by arrows. There is a slight conformational difference in the position of residue $M174^{5.35}$ between the two structures, as shown by the cryo-EM maps. In the 8X16 structure the position of $M174^{5.35}$ occludes the $N^6$-iodobenzyl group from a cryptic pocket created by ECL2 and TM5.

**Table S1. Cryo-EM data collection, refinement, and validation statistics**

| | A3AR-DNGi1-scFv16-adenosine | A3AR-mGsi-Nb35-Piclidenoson | A3ARBRIL-BAG2-Nb-LUF7602 |
|---|---|---|---|
| **Data Collection** | | | |
| PDB code | 9EBH | 9EBI | 9EHS |
| EMD code (Consensus)<br>EMD code (Receptor Focus)<br>EMD code (Composite) | 47879<br>47994<br>N/A | 47880<br>47998<br>N/A | 48063<br>48064<br>48065 |
| Micrographs | 8686 | 6526 | 6421 |
| Electron Dose ($e^-/A^2$) | 60 | 60 | 60 |
| Voltage (kV) | 200 | 300 | 300 |
| Pixel size (Å) | 0.99 and 1.03 | 0.82 | 0.65 |
| Movie frames | 50 | 60 | 60 |
| Defocus range (µm) | 0.5 – 1.5 | 0.5 – 1.5 | 0.5 – 1.5 |
| **Refinement** | | | |
| Symmetry imposed | C1 | C1 | C1 |
| Particles (final map) | 325k | 590k | 328K |
| Resolution @0.143 FSC (Å)*<br>Complex<br>Receptor local refinement | <br>3.05<br>3.44 | <br>2.48 | <br>2.65<br>3.34 |
| $CC_{map–model}$ (volume) | 0.71 | 0.85 | 0.78 |
| **Model Quality** | | | |
| R.M.S. deviations | | | |
| Bond length (Å) | 0.003 | 0.005 | 0.004 |
| Bond angles (°) | 0.716 | 0.795 | 0.775 |
| Ramachandran | | | |
| Favoured (%) | 98.89 | 99.0 | 99.02 |
| Outliers (%) | 0 | 0 | 0 |
| Rotamer outliers (%) | 0 | 0.35 | 0 |
| C-beta deviations (%) | 0 | 0 | 0 |
| Clashscore | 4.02 | 4.90 | 2.86 |
| MolProbity score | 1.19 | 1.26 | 1.08 |

**Table S2. Potency and $E_{max}$ values from the $G\alpha_{i1}$ activation assay (TruPath)**

| A3 Adenosine Receptor | Adenosine pEC$_{50}$ (n) | Piclidenoson pEC$_{50}$ (n) | NECA pEC$_{50}$ (n) |
|---|---|---|---|
| **WT** | 6.68 ± 0.06 | 7.86 ± 0.13 | 7.38 ± 0.15 |
| **Y15A$^{1.35}$** | 4.16 ± 0.36 **** | 6.50 ± 0.42 *** | 5.02 ± 0.02 **** |
| **S73A$^{2.65}$** | 5.90 ± 0.11 * | 7.38 ± 0.03 | 6.73 ± 0.14 ** |
| **T94A$^{3.36}$** | 5.01 ± 0.13 **** | 6.76 ± 0.16 ** | 5.47 ± 0.10 **** |
| **H95A$^{3.37}$** | 3.57 ± 0.06 **** | N.R. | 5.14 ± 0.07 **** |
| **H95F$^{3.37}$** | N.R. | N.R. | N.R. |
| **V169E$^{45.53}$** | 6.64 ± 0.17 | 7.94 ± 0.01 | 7.40 ± 0.05 |
| **M174A$^{5.35}$** | 5.97 ± 0.12 * | 8.16 ± 0.06 | 6.85 ± 0.06 * |
| **N250$^{6.55}$** | N.R. | N.R. | N.R. |
| **Y265A$^{7.36}$** | 6.01 ± 0.12 * | 7.44 ± 0.07 | 7.05 ± 0.09 |
| **S271A$^{7.42}$** | 3.19 ± 0.10 **** | N.R. | 4.34 ± 0.15 **** |
| **H272$^{7.43}$** | N.R. | N.R. | N.R. |
| **A3 Adenosine Receptor** | **Adenosine** %E$_{max}$ (n) | **Piclidenoson** %E$_{max}$ (n) | **NECA** %E$_{max}$ (n) |
| **WT** | 6.68 ± 0.06 | 7.86 ± 0.13 | 7.38 ± 0.15 |
| **Y15A$^{1.35}$** | 4.16 ± 0.36 **** | 6.50 ± 0.42 *** | 5.02 ± 0.02 **** |
| **S73A$^{2.65}$** | 5.90 ± 0.11 * | 7.38 ± 0.03 | 6.73 ± 0.14 ** |
| **T94A$^{3.36}$** | 5.01 ± 0.13 **** | 6.76 ± 0.16 ** | 5.47 ± 0.10 **** |
| **H95A$^{3.37}$** | 3.57 ± 0.06 **** | N.R. | 5.14 ± 0.07 **** |
| **H95F$^{3.37}$** | N.R. | N.R. | N.R. |
| **V169E$^{45.53}$** | 6.64 ± 0.17 | 7.94 ± 0.01 | 7.40 ± 0.05 |
| **M174A$^{5.35}$** | 5.97 ± 0.12 * | 8.16 ± 0.06 | 6.85 ± 0.06 * |
| **N250$^{6.55}$** | N.R. | N.R. | N.R. |
| **Y265A$^{7.36}$** | 6.01 ± 0.12 * | 7.44 ± 0.07 | 7.05 ± 0.09 |
| **S271A$^{7.42}$** | 3.19 ± 0.10 **** | N.R. | 4.34 ± 0.15 **** |
| **H272$^{7.43}$** | N.R. | N.R. | N.R. |

Mean ± SEM pEC$_{50}$ and % maximum response (Emax) values for adenosine, NECA, and Piclidenoson from HEK293A cells transiently expressing WT A$_3$AR and single residue mutants. N.R. denotes mutants which a measurable response could not be detected. Values are calculated from three independent experiments conducted in duplicate. Statistically significant differences compared to WT A$_3$AR were calculated using a one-way ANOVA with a Dunnett's multiple comparison test. P values: *P ≤ 0.05, **P ≤ 0.01, ***P ≤ 0.001, and ****P ≤ 0.0001.
